# Hepatic Yap1 activates systemic catabolism and muscle loss during organ repair: evidence for a liver-derived common mechanism with cancer cachexia

**DOI:** 10.1101/2025.04.01.646698

**Authors:** Tewfik Hamidi, Yanlin Jiang, Tyler Robinson, Andris Kronbergs, Ernie Au, Joseph E. Rupert, Xiaoling Zhong, Siangsin Lal, Tiffany Liang, Alexander G. Robling, April Hoggatt, Robson F. Carvalho, Sarah S. Cury, Clark D. Wells, Teresa A. Zimmers, Leonidas G. Koniaris

## Abstract

Recovery from critical injury concomitant with restoration of functional organ mass invokes a systemic catabolic response leading to muscle and fat loss, known as cachexia. We interrogated this process using mouse models of organ repair, including liver regeneration after hepatectomy, Yap1-mediated hepatomegaly, and pneumonectomy. Both hepatectomy and Yap1 induced systemic catabolism. Muscle and adipose wasting scaled with degree of liver growth, with 10-25% reduction of muscle mass and 50-80% of fat mass. In contrast, non-regenerative lung injury did not induce tissue wasting. Liver growth elevated resting energy expenditure. Tracer studies demonstrated redistribution of muscle-derived cholesterol and amino acids to regenerating liver. Gene expression changes in livers and muscles showed high concordance between the liver growth models and cancer cachexia models, including pancreatic adenocarcinoma cachexia. We propose that cachexia is a normal and essential reparative process in organ repair and regeneration and further, that cancer cachexia is a pathological exacerbation of an adaptive process mediated by activation of Yap1 in the liver.

## Introduction

Surviving critical organ injury, sepsis or surgery requires an essential catabolic recovery period that typically extends from days to weeks ^1^. This catabolism involves the breakdown of existing molecules into smaller units that are either oxidized to release energy or used in other anabolic reactions. The catabolic response is systemic, activates a breakdown of fat and skeletal muscle during organ repair, and resolves with recovery. A rapid loss of muscle in long-bone fracture was first reported by Cuthbertson in 1930 and was termed “ebb and flow” ^2^. This process has subsequently been termed “hypermetabolism” or “the adrenergic-corticoid phase” ^3,4^. Work by Rhoads and others found that this catabolic response, rather than nutritional intake, drives repair and regeneration of tissues following injury ^5^. Serious injuries including major trauma, liver resection, and burns can require catabolic responses over days to weeks to fully recover. Interestingly, metabolic reprogramming constitutes one of the major hallmarks of cancer^6^. Similar to critically injury patients, a large majority of patients with advanced-stage cancer show a severe catabolic syndrome associated with muscle and fat loss, also termed cancer cachexia ^7^. Nutritional support to overcome cancer cachexia has been so far challenging ^8^. Although optimizing preoperative and postoperative nutritional support in these patients improves surgical outcomes, it does not prevent tissue catabolism. Conversely, an impaired catabolic response may be associated with increased morbidity and mortality. Although current literature has focused on pathological persistence of the catabolic response and energy expenditure following injury, particularly after burns ^9^, acute catabolism is essential to survive major injuries and its underlying mechanisms remain poorly understood.

Organ damage induces several signaling pathways, including Il-6/Jak/Stat3, Tnfα, Hippo/Yap1/Taz, that are essential for the initiation and execution of the reparative processes^10^. The hippo pathway targets, Yap1 and Taz, mainly drive the proliferative arm of the reparative process ^9,11–14^. Yap1/Taz transcriptional co-activators play key roles in organ size control, as well as during organ injury repair including liver regeneration ^15^. The actions of Yap1 and Taz on the overall body metabolism remain an active area of investigation. For instance, it is poorly understood how organ repair and tissue-focused anabolic signals induce systemic catabolic response.

Herein, we identify Yap1 induced liver proliferation as a driver of the catabolic response through the production of liver derived serum factors. Both liver regeneration post-hepatectomy and Yap1-induced liver expansion, hereafter termed liver growth mouse models, induce a systemic catabolic response in mice involving increased resting energy expenditure (REE) and associated with significant loss of muscle and fat mass. Furthermore, Yap1 activation in the liver results in upregulated skeletal muscle fatty acid oxidation, branched chain amino acids (BCAA) catabolism, lipid transport signaling, as well as accumulation of oxidized lipids and the release of free BCAA into the circulation. During liver growth-induced catabolic process, we showed that muscle-derived amino acids and lipids relocate to the proliferating liver, suggesting the use of muscle and fat substrates to support liver proliferation. Thus, Yap1 is demonstrated to promote organ regeneration after injury through the modulation of whole-body metabolism. Finally, gene expression changes in liver and skeletal muscle from liver growth mouse models predicted numerous alterations in metabolic signaling pathways that recapitulate effects observed in cancer cachexia mouse models. Our data suggest that cancer cachexia may represent a pathological activation of liver reparative response.

## Results

### Post-hepatectomy liver regeneration induces body weight loss, muscle and fat loss in mice

Organ injury induces a stress response associated with damage repair. To investigate the catabolic phenotype following liver resection, 70% partial hepatectomies (PHx) were performed on C57BL6 male mice ^16^ (Supplementary Fig. 1A). Body composition was monitored over time. Mice were healthy and recovered 78% of liver mass on average by 7 days post-surgery (Fig. 1A). PHx-mice showed on average a 20% decrease in body mass and 23.3% reduction in carcass mass at day 7 versus sham surgery mice (Fig. 1B). By day 7, PHx-mice showed a 24% decrease in quadriceps and gastrocnemius mass and a 26 % loss of tibialis anterior (TA) mass (Fig. 1C). Consistent with gastrocnemius mass loss, we found a 26.7% reduction in fiber cross sectional area (CSA) (Fig. 1D). Compared to sham mice, a higher frequency of thin fibers (CSA from 1000 to 2000 µm^2^) was observed in PHx-mice. Conversely, PHx-mice showed fewer thick fibers (CSA from 2400 to 4200 µm^2^) compared to sham mice (Fig. 1E). Functional analysis showed that post-PHx mice had a 28.1% and 21% reduction in grip strength compared to sham mice at day 7 and day 14 respectively, demonstrating a loss in muscle function post-PHx (Supplementary Fig. 1B). Histological analysis of epidydimal white adipose tissue (eWAT) showed a clear reduction in adipocyte size in PHx-mice (Fig. 1F). Consistently, eWAT mass was reduced by 46.4% and 79.4% at days 3 and 7 post-PHx respectively (Fig. 1G). To determine the relative contribution of surgical trauma versus the regenerative response, we performed left pneumonectomies (Pn), removing approximately 45% of total lung volume. Pneumonectomy, a non-regenerative injury, did not cause body weight loss (not shown) nor skeletal muscle loss (Supplementary Fig. 1C). Liver regeneration involves compensatory hyperplasia by a subset of remaining hepatocytes until the entire organ mass is recovered. Histologically, both increased proliferation and cellular enlargement are observed ^16,17^. Levels of the cell proliferation marker Pcna were elevated at day 2, 3 and 7 in PHx-livers, consistent with other reports (Fig. 1H). In parallel, Yap1, Taz and phosphor-Stat3 protein levels increased at each of these days, demonstrating active reparative processes post-PHx. This was accompanied by reduced levels of phosphorylated Yap1, demonstrating its activation and nuclear localization (Fig. 1H). Gene expression of Yap1/Taz target genes *Ctgf*, *Cyr61* and *Il-6* were significantly increased 24 hours after PHx (Fig. 1I). Given the significant loss in skeletal muscle mass after PHx, we analyzed the common muscle atrophy signaling pathways^18^. We found the fork head box (FoxO) family transcription factors FoxO1 and FoxO3a elevated in gastrocnemius from PHx-mice (Fig. 1J). Muscles also showed elevated Akt signaling (p-Akt/Akt), autophagy markers (LC3II/LC3I ratio) and ubiquitination (Fig. 1J). Moreover, an upregulation of muscle atrophy and autophagy related genes *Fbxo32* (Atrogin1), *Trim63* (Murf1), *Bnip3* and *Bnip3l* was observed at days 1, 2 and 7 post-PHx (Fig. 1K). Taken together, liver regeneration and repair cause a severe body weight loss, associated with decreased fat and skeletal muscle mass. Muscle loss is associated with the activation of Akt, FoxO pathways and the increased expression of muscle atrophy factors and the subsequent activation of proteolysis hallmarks, including ubiquitination and autophagy.

**Fig. 1:**
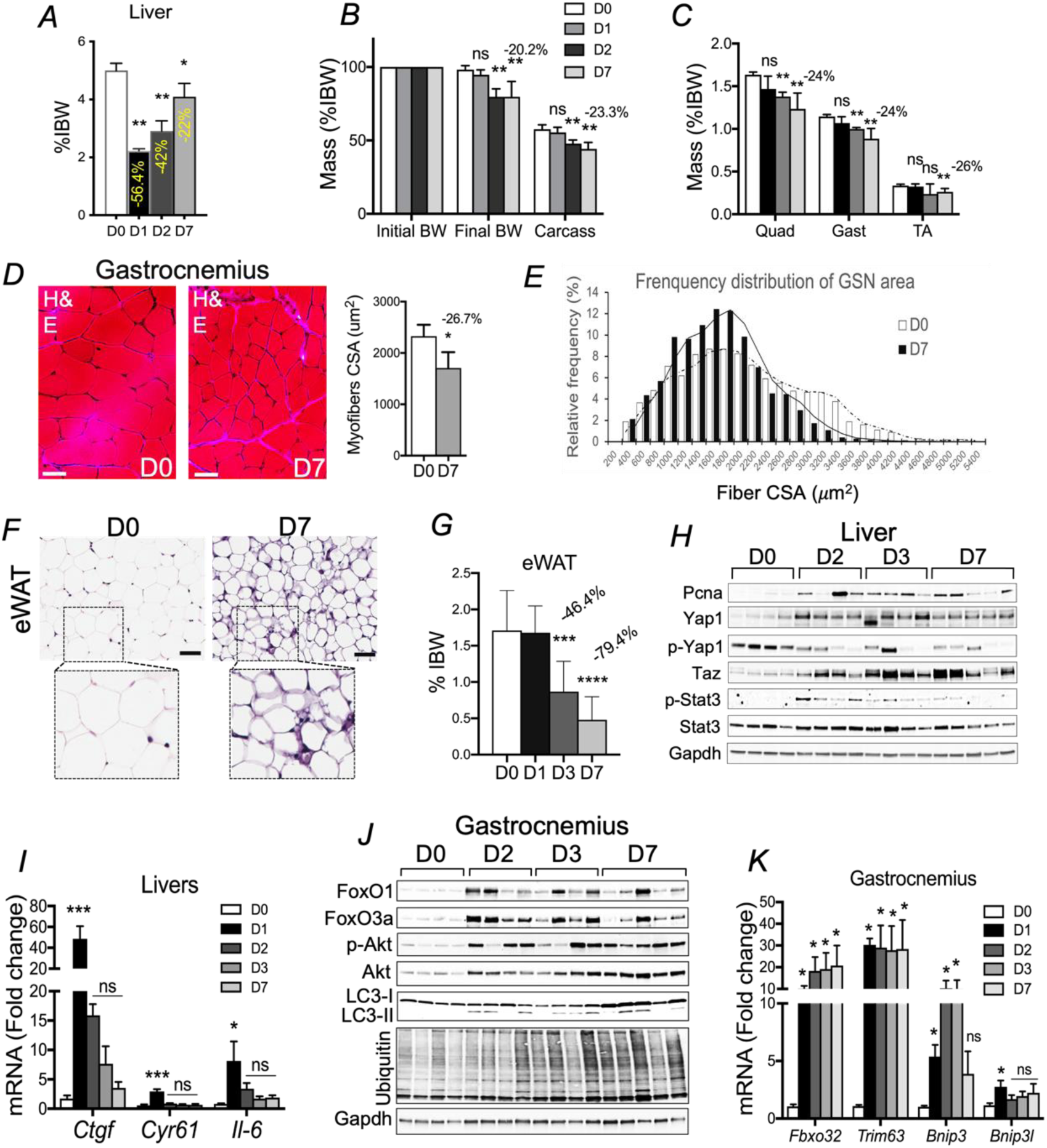
Post-hepatectomy liver regeneration induces body weight loss, muscle and fat loss in mice. A) Liver mass shown as percentage of initial body weight (%IBW) at day0, day1, day2 and day7 post-PHx. n=5 mice per condition, *P<0.05, **P<0.01, One-Way ANOVA. B) Final body weight, carcass weight at day0, day1, day2, day7. n=5, **P<0.01, ns: not significant, One-Way ANOVA. C) Skeletal muscle mass at day0, day1, day2, day7. n=5, **P<0.01, ns: not significant, One-Way ANOVA. D) H&E staining of gastrocnemius cross section at day0 versus day7, and quantification of myofibers cross section area (CSA). n=5 mice per condition analyzed. *P<0.05, unpaired t-test. Scale bar= 50 microns. E) Frequency distribution of gastrocnemius CSA at day0, day7. CSAs were measured from: n=5 day0 mice, n=7 day7 mice. F) Representative H&E staining of epididymal white tissue adipose (eWAT) at day0, day7 post-PHx. Scale bar = 50 microns. G) Fat mass (eWAT) at day0, day1, day3, day7. n=5 mice per condition, ***P<0.001, ****P<0.0001, One-Way ANOVA. H) Western blot analysis of Pcna, p-Yap1, Yap1, Taz, Stat3 and p-Stat3 in livers at day0, day2, day3, day7 post-PHx. I) Gene expression of *Ctgf*, *Cyr61* and *Il-6* in post-PHx livers. n=5 mice per time point, \**P*<0.05, \*\*\**P*<0.001, One-Way ANOVA. J) Western blot analysis of FoxO1, FoxO3a, p-Akt, Akt, LC3 and ubiquitin in gastrocnemius at day0, day2, day3, day7. K) Gene expression of *Trim63, Fbxo32*, *Bnip3, Bnip3l* in gastrocnemius. n=5 mice per time point, \**P*<0.05, One-Way ANOVA.

### Yap1-induced liver growth causes body weight loss, muscle and fat loss in mice

Hepatectomy activated Yap1/Taz and increased the expression of their target genes in the liver (Fig. 1H-I). The possibility that hepatic Yap1 activation would be sufficient to cause a reduction in body, muscle and fat mass was then determined. For this purpose, mice were infected with an adeno-associated virus carrying the constitutively active *Yap1^S5A^* mutant under the control of the liver specific TBG promoter ^19^, hereafter termed AAV-Yap1. An approximate doubling of liver size was observed in AAV-Yap1 mice compared to AAV-GFP after 7-and 14-days (Fig. 2A, B). AAV-Yap1 mice showed no body weight (FBW) loss at day7, most likely due to the increased liver mass (+127%) that could have unmasked the combination of fat and muscle mass loss (Fig. 2C). However, final body weight was significantly in AAV-Yap1 mice at 14 days, illustrating a severe fat and muscle loss reducing the total body weight even in presence of a 99% increase in liver mass due to active Yap1 expression. AAV-Yap1 mice lost from 8.8 to 13% of muscle mass at day7 and 10% to 14% at day 14 (Fig. 2D). Additionally, a dramatic 56.4% reduction in eWAT mass is shown at day 7 (Fig. 2E). Consistently, the average muscle CSA in AAV-Yap1 mice was reduced by 6.3% at day 7 (Fig. 2F). Assessment of muscle strength, using grip test and hanging test determined that muscle loss was associated with decreased muscle strength (Supplementary Fig. 2A). Analysis of AAV-Yap1 livers confirmed Yap1 hepatic and increased levels of Pcna levels at 7 and 14 days (Fig. 2G). Elevated Pcna and Ki67 levels were also detected in AAV-Yap1 liver sections, demonstrating proliferative liver tissues (Fig. 2J). Activation of Yap1 at days 7 and 14 was further confirmed by increased transcript levels of *Il-6*, *Ctgf* and *Cyr61* (Fig. 2H). Since Il-6/Jak/Stat3 signaling promotes regeneration in injured livers ^20^, we monitored Il-6 transcript and protein levels in concert with phospho-Stat3 levels. Hepatic Il-6 mRNA and protein levels were increased in livers at day7-and day14 post AAV-Yap1 infection (Fig. 2G-I). Moreover, phospho-Stat3 levels were elevated in these livers (Fig. 2G-H). Hepatic Il-6/Jak/Stat3 signaling is therefore induced as a result of hepatic Yap1 activation. Consistent with the severe loss in skeletal muscle mass and function observed in AAV-Yap1 mice, we found that ubiquitination levels, as well as FoxO3 levels, were significantly increased in AAV-Yap1 muscle (Fig. 2K-L). In a complementary approach, we examined liver growth and muscle wasting in the doxycycline responsive *ApoE-rtTA-Yap1^S5A^* transgenic model (Fig. 2M) ^21^. For simplicity, water fed *ApoE-rtTA-Yap1^S5A^*mice are termed Yap1^OFF^, whereas *ApoE-rtTA-Yap1^S5A^*mice fer with doxycycline will be termed Yap1^ON^. We confirmed that hepatic expression of Yap1 in response to doxycycline exposure was liver specific (Supplementary Fig. 2B). Yap1^ON^ livers grew in mass by 260 % and 374 % at day 7 and 14, respectively (Fig. 2N). In parallel, Yap1^ON^ mice experienced a 60.3% and 90.7% loss of eWAT mass at day 7 and 14 and skeletal muscle mass was reduced by 25-30% at day 14 (Fig. 2N). Altogether, data show that, similarly to liver regeneration model, activation of hepatic Yap1 causes body weight loss, muscle and fat loss in mice. Yap1 activation induces the Il-6/Jak/Stat3 signaling in the liver and skeletal muscle loss through FoxO3 mediated activation of proteasomal degradation.

**Fig. 2:**
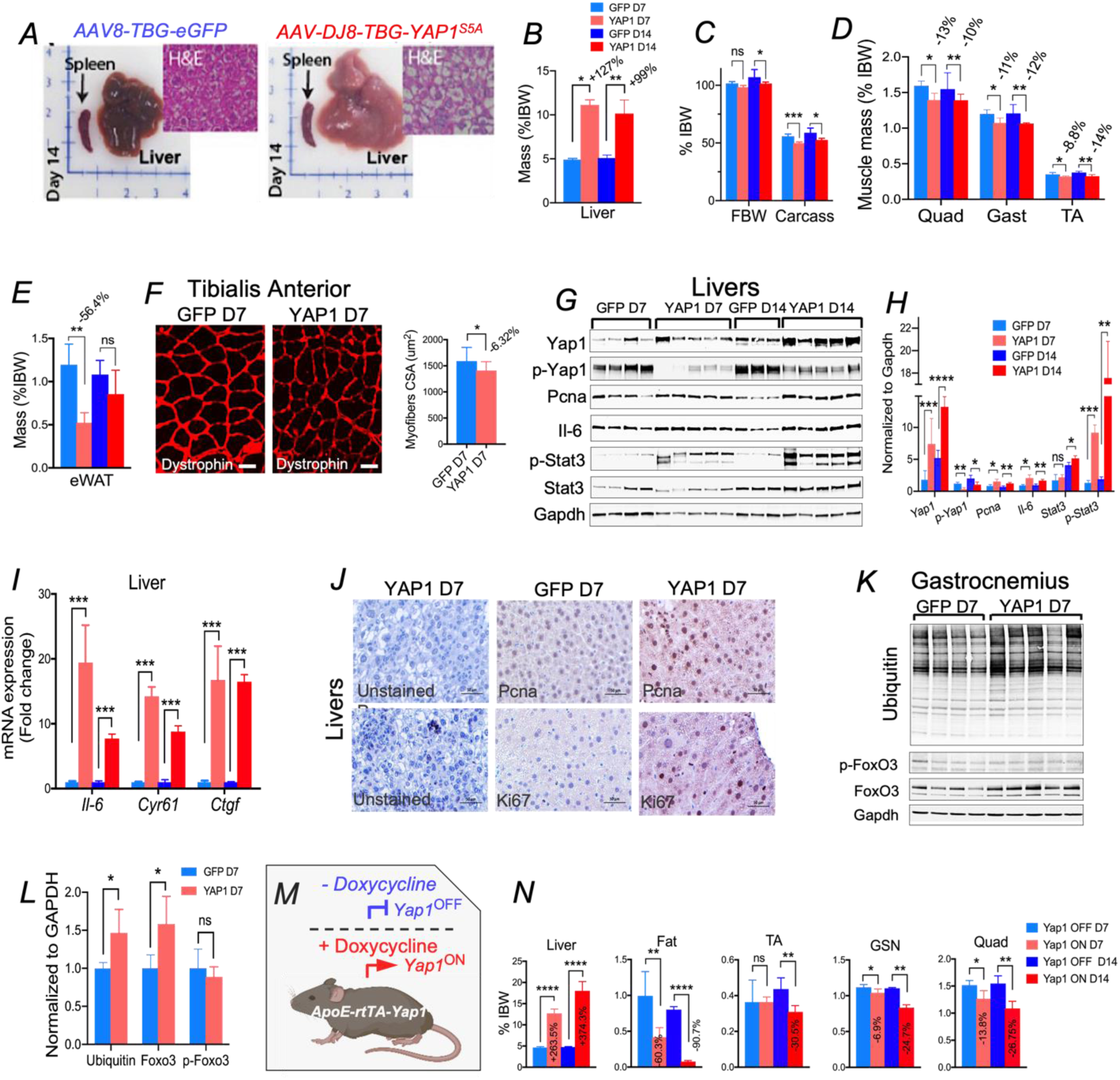
Yap1-induced liver growth causes body weight loss, muscle and fat loss in mice. A) Hepatic expression of active Yap1^S5A^ induces hepatomegaly. *AAV8-TBG-eGFP* (GFP) and *AAV-DJ8-TBG-Yap1^S5A^* (Yap1) were injected into mice. AAV-GFP: n=7 mice, AAV-Yap1: n=10 mice. B) Liver mass measured at day7 and day14 post-AAV injection. \**P*<0.05, \*\**P*<0.01. C) Final body weight (FBW) and carcass weight at day7, day14 post-infection. *P<0.05, ***P<0.001. D) Skeletal muscle weight at day7, day14. *P<0.05, **P<0.01. E) eWAT mass measured at day7, day14. **P<0.01, ns: not significant. F) Representative images of TA sections stained with dystrophin (Red) along with quantification of CSA in both conditions. n=5 mice per condition, \**P*<0.05. G) Western blot analysis of Pcna, Il-6, p-Sta3/Stat3, p-Yap1/Yap1 in livers at day7, day14 post-AAV infection. H) Band intensities in (G) were quantified and normalized to Gapdh levels. *P<0.05, **P<0.01, ***P<0.001, One-way ANOVA. I) Gene expression changes of *Ctgf*, *Il-6* and *Cyr61* in the liver at day7, day14 post-injection. ***P<0.001, One-way ANOVA. J) Immunohistochemistry staining for Pcna, Ki67 at day7 post-infection. Scale bars= 50 microns. K) Western blot analysis of muscle Ubiquitin, Foxo3, p-Foxo3 at day7 post-injection. L) Band intensities in (K) were quantified. *P<0.05, unpaired t-test. M) Hepatic expression of active Yap1^S5A^ using the doxycycline responsive *ApoE-rtTA-Yap1^S5A^*mouse model. [*ApoE-rtTA-Yap1^S5A^* minus doxycycline = Yap1^OFF^], [*ApoE-rtTA-Yap1^S5A^* plus doxycycline = Yap1^ON^]. N) Liver, fat (eWAT) and muscle mass measured at day7 and day14 after doxycycline administration. Yap1^OFF^: n=5 mice, Yap1^ON^: n=8 mice. *P<0.05, **P<0.01, ****P<0.0001.

### Yap1-mediated hepatomegaly induces body weight loss and muscle loss independently of Il-6 and Ctgf

Interleukin-6 plays a major role in mediating the catabolic response to trauma and severe illness ^22–24^. We previously showed that systemic administration of Il-6 induces hepatocyte proliferation and hepatomegaly ^25^. Furthermore, in tumorigenic contexts, including liver cancer, hepatic Yap1 controls Il-6 transcription in the liver ^26^ ^27^. To test whether Il-6-induced hepatocyte proliferation may cause skeletal muscle loss, mice were infected with AAV-CAG-mIl-6. At day7, mice overexpressing mIl-6 showed a 42 % increase in liver mass compared to control mice (Supplementary Fig. 3A). mIl-6 mice showed a reduction in TA mass, gastrocnemius and quadriceps mass of 28.5%, 20% and 32.4% respectively, whereas the heart mass was unaffected (Supplementary Fig. 3B). This observation raises the possibility that Yap1-induced *Il-6* expression promotes skeletal muscle atrophy during liver growth. To test this possibility, *Il-6* deficient mice were infected with AAV-Yap1 similar to (Fig. 2A). As expected, AAV-Yap1 induced hepatomegaly, body weight loss and muscle loss in wild-type mice (Supplementary Fig. 3C-F). However, Yap1 induced similar liver growth and reduced body weight, carcass weight loss in *Il-6* deficient mice (Supplementary Fig. 3C-E). Yap1-associated skeletal muscle loss was similar between *wild-type* and *Il-6* deficient animals even 8 weeks after AAV injection (Supplementary Fig. 3F). This data indicates that Il-6 is essential to induce liver growth, but not necessary for Yap1-induced muscle loss upon liver injury and growth. We next tested the role of another Yap1 target gene (*Ctgf*) in liver growth and muscle loss. Connective tissue growth factor is secreted in response to liver damage to regulate hepatocyte differentiation ^28,29^. While AAV-Ctgf mice showed sustained overexpression of hepatic Ctgf for 24 days (Supplementary Fig. 4A), this did not produce hepatomegaly. Only a modest increase in liver mass was observed 56 days post infection (Supplementary Fig. 4B-C). Concordantly, *Ctgf* overexpression did not affect body weight or skeletal muscle weight by 56 days post-infection (Supplementary Fig. 4D, E). Taken together, we show that neither Il-6 nor Ctgf were found essential for the effects of Yap1 on liver growth and muscle wasting. We therefore sought to further investigate the liver/muscle crosstalk mechanisms that promote muscle loss in the liver growth models.

### Liver-derived circulating serum factors during liver growth inhibit myogenesis and induce myotube atrophy *in vitro*

Many cellular functions, especially metabolism and organ size are integrated by crosstalk between the liver and other tissues ^30,31^. We therefore tested whether regenerating liver releases signaling factor(s) that enter the circulation and may induce skeletal muscle atrophy. Blood plasma collected from sham or PHx mice was added to differentiated C_2_C_12_ myotubes (Fig. 3A). Myotube diameter and nuclei number per myotube were measured to assess myotube atrophy and myogenesis respectively. Myotubes treated with plasma from 48hr post-PHx-mice showed a 12% reduction in diameter and a 27% less nuclei per fiber (Fig. 3A). However, myotubes treated with plasma from 24hr post-PHx-mice did not show significant changes. These findings indicate that regenerating livers promote the secretion of factor(s) into the blood that induce muscle atrophy. Since blood plasma was used here, it was unclear if the liver is the direct source of these factor(s). To test this, post-PHx livers were used for C_2_C_12_ media conditioning (Fig. 3B). Myotubes treated with post-PHx liver conditioned media (CM) showed a 9.7% reduction in their diameter and 28.6% fewer nuclei per myotube, demonstrating that post-PHx liver factors inhibit myogenesis and induce muscle atrophy in vitro. Similarly, blood plasma collected from Yap1^ON^ mice (*ApoE-rtTA-Yap1^S5A^*+ doxycycline) reduced myotubes diameter by 16.55% and reduced the number of nuclei per fiber by 40% (Fig. 3C). Collectively, our data show that factors secreted directly from regenerating livers are sufficient to induce atrophy and impair myogenesis in cultured C_2_C_12_ myotubes. Moreover, our data also indicate that hepatic Yap1 generates a secretory signature that promotes muscle atrophy.

**Fig. 3:**
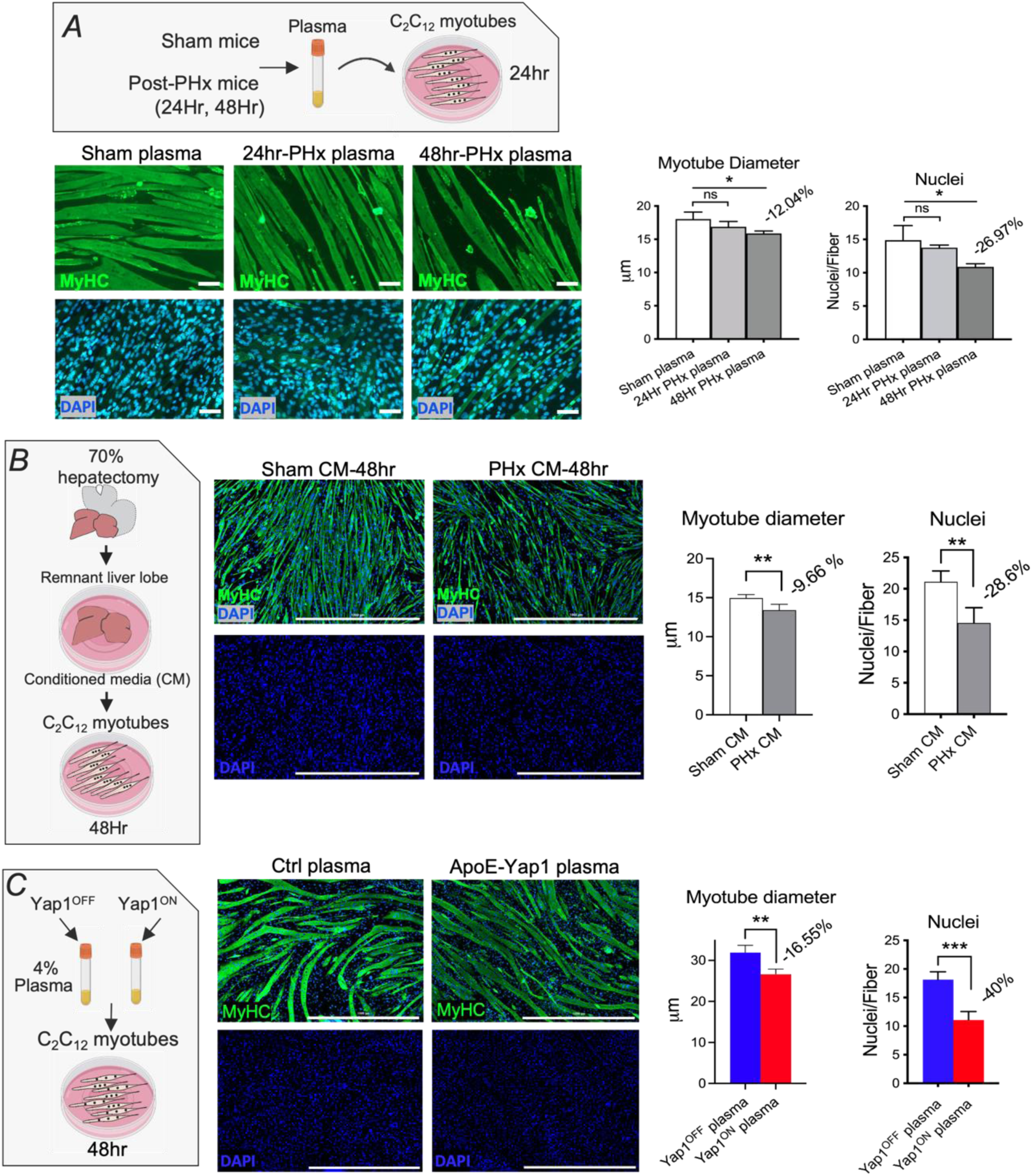
Liver-derived circulating serum factors during liver growth inhibit myogenesis and induce myotube atrophy *in vitro*. A) Top panel shows experimental design. Blood plasma was collected from 24 hr and 48 hr post-PHx-mice. Media + 4% plasma was then added to C_2_C_12_ myotubes to test plasma activity for 24 hrs. Bottom left panel shows images of MyHC/DAPI stained myotubes after incubation with plasmas. Scale bars = 50 microns. Right panel shows quantification of myotubes diameter and nuclei number per myofiber in each condition. n=2 mice per condition, 3 C_2_C_12_ wells tested for each mouse. *P<0.05, One-way ANOVA. B) Left panel shows experimental design. C_2_C_12_ myotubes were incubated with Sham-or PHx-liver conditioned media (CM) for 48 hrs. Middle panel shows images of MyHC/DAPI stained myotubes after incubation with Sham-CM, or PHx-CM. Scale bar= 1000 microns. Right panel shows quantification of myotubes diameter and nuclei number per myofiber. n=2 sham mice and 3 PHx-mice tested. 3 C_2_C_12_ wells tested for each mouse. **P<0.01, One-way ANOVA. C) Left panel, plasma was collected from *Wt* or *ApoE-rtTA-Yap1^S5A^* mice fed with doxycycline for 14 days. Plasma was then added at 4% in the C_2_C_12_ differentiation media for 48 hrs. Middle panel shows images of MyHC/DAPI stained myotubes in each condition. Scale bar = 1000 microns. Right panel, myotubes diameter quantification and nuclei per myofiber in each condition. n=3 mice tested per condition. n=2 C_2_C_12_ wells tested for each mouse. **P<0.01, ***P<0.001, One-way ANOVA.

### Liver growth and repair increase resting energy expenditure and fat metabolism in mice

Hypermetabolism is often observed in critically ill patients and correlates with a severe weight loss ^32–35^. The severe body weight loss post-PHx and Yap1-induced hepatomegaly spurred an investigation into the metabolic changes in these mouse models. The respiratory quotient (RQ) measures the volume of carbon dioxide released over the volume of dioxygen absorbed (VCO_2_/VO_2_). RQ is influenced proportionately by the relative oxidation of fat by (0.7), protein by (0.8), and carbohydrate by (1.0) ^36,37^. Post-PHx mice were housed in metabolic cages that monitor O_2_ consumption, CO_2_ release, food/water intake and activity for up to 5 days. Compared to sham group, PHx-mice were less active and consequently consumed less O_2_ and exhaled less CO_2_, presumably due to trauma from surgery (Fig. 4 A-C). We found a reduced RQ in PHx-mice (∼0.7) at days 2, 3 and 4 compared to sham animals (∼0.8), indicating that PHx-mice increased the use of fat as an energy substate (Fig. 4D), consistent with the rapid fat loss observed post-PHx (Fig. 1F-G). We observed a significant increase in resting energy expenditure (REE) early post-PHx when normalized to activity, demonstrating a rapid induction of hypermetabolism in PHx-mice (Fig. 4E), presumably due to the metabolic needs for liver repair. PHx-mice also showed reduced food and water intake during the first 48 hrs after partial hepatectomy (Fig. 4F-G), most likely due to surgical trauma. Taken together, we show that liver regeneration induces a hypermetabolic state, associated with increased fat use as an energy substrate. We next tested the metabolic profile of Yap1^ON^ mice. Due to absence of surgical trauma, Yap1^ON^ and Yap1^OFF^ mice showed similar activity levels (Fig. 4H), as well as similar food and water intake (Fig. 4M-N). Overall, O_2_ consumption and generation of CO_2_ were elevated in Yap1^ON^ mice compared to the Yap1^OFF^ group (Fig. 4I, J). Yap1^ON^ mice RQ was slightly but significantly decreased at days 3 and 4, demonstrating increased use of fat as an energy substrate (Fig. 4K). REE also showed an overall increase in metabolism in Yap1^ON^ mice at days 3, 4, and 5 after exposure to doxycycline (Fig. 4L). Taken together, metabolic profiling shows that liver regeneration and growth trigger a systemic catabolic state, including elevated resting energy expenditure and a significant increase in fat use as an energy substrate.

**Fig. 4:**
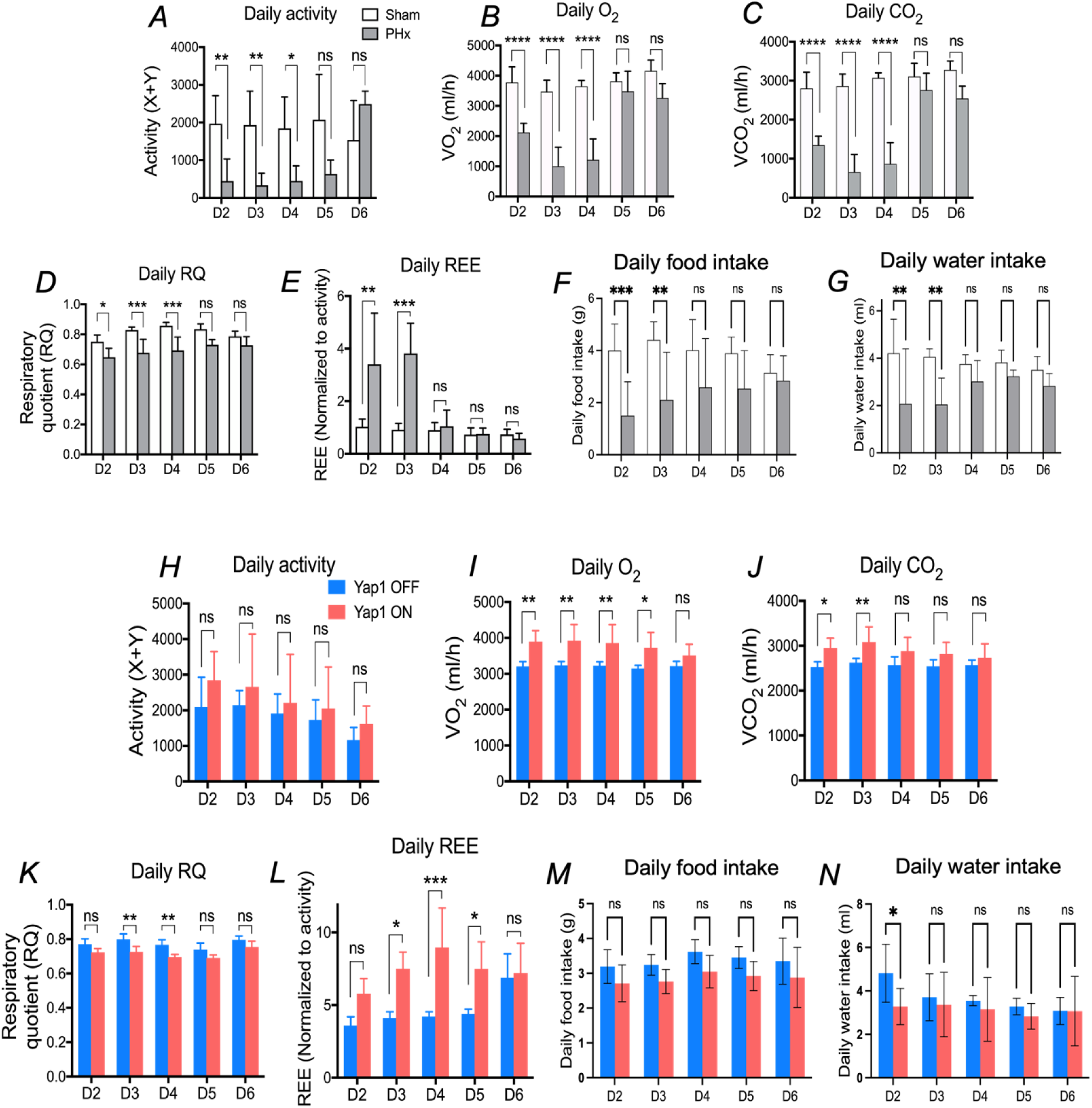
Liver growth and repair increase resting energy expenditure and fat metabolism in mice. A) Daily activity measured from day2 to day6 post-PHx. n=5 sham mice, n=9 PHx-mice. *P<0.05, **P<0.01, Two-way ANOVA. B) Daily consumed dioxygen volume post-PHx. \*\*\*\**P*<0.0001, Two-way-ANOVA. C) Daily exhaled carbon dioxide volume post-PHx. \*\*\*\**P*<0.0001, Two-way ANOVA. D) Daily respiratory quotient (RQ) post-PHx. \**P*<0.05, \*\*\**P*<0.001, Two-way ANOVA. E) Daily resting energy expenditure (REE) post-PHx, normalized to mice activity in (A). **P<0.01, \*\*\**P*<0.001, Two-way ANOVA. F, G) Daily food and water intake in sham and PHx mice. \*\**P*<0.01, \*\*\**P*<0.001, Two-way ANOVA. H) Daily activity of Yap1^OFF^ and Yap1^ON^ mice measured from day2 to day6 post-doxycycline administration. n=5 Yap1^OFF^, n=9 Yap1^ON^ mice. Two-way ANOVA. I) Daily consumed dioxygen volume of Yap1^OFF^ and Yap1^ON^ mice. \**P*<0.05, \*\**P*<0.01, Two-way ANOVA. J) Daily exhaled carbon dioxide volume of Yap1^OFF^ and Yap1^ON^ mice. \**P*<0.05, \*\**P*<0.01, one-way ANOVA. K) Daily RQ in Yap1^OFF^ and Yap1^ON^ mice. \*\**P*<0.01, Two-way ANOVA. L) Daily normalized REE. \**P*<0.05, \*\*\**P*<0.001, Two-way ANOVA. M, N) Daily food and water intake in Yap1^OFF^ and Yap1^ON^ mice and PHx mice. \**P*<0.05, Two-way ANOVA.

### Liver and skeletal muscle gene expression in the liver growth mouse models

We next examined hepatic and muscle gene expression changes in both PHx and AAV-Yap1 mice. Post-PHx livers and quadriceps showed 4080 and 6177 differentially expressed genes (DEGs) respectively (Fig. 5A), whereas AAV-Yap1 livers and quadriceps resulted in 6616 and 1124 differentially expressed genes (Fig. 5B). Canonical pathway analysis of hepatic gene expression in both models revealed that most significantly affected pathways are related to cell proliferation, integrins, extracellular matrix organization, inflammatory response and numerous metabolic functions (Fig. 5C-D, left panels, supplementary table 1), consistent with previously reported pathways associated with liver growth and response to damage ^38,39^. In parallel, quadriceps from both models showed an enrichment of canonical pathways associated with muscle atrophy, such as protein catabolic process, proteasome, FoxO signaling, autophagy, complement and coagulation cascade, NFκB, BCAA catabolism and fatty acid oxidation (Fig. 5C-D, right panels, supplementary table 1). Together, gene expression analysis confirmed that liver growth in response to injury or Yap1 activation induces skeletal muscle atrophy via the upregulation of signaling pathways promoting proteolysis, autophagy and lipid oxidation. Elevated BCAA catabolism and fatty acid beta oxidation gene signatures in skeletal muscle suggest the breakdown of muscle-derived large molecules (proteins and lipids) to increase the availability of anabolic metabolites to support liver growth. Gene set enrichment analysis (GSEA) confirmed that BCAA catabolism and mitochondrial fatty acid oxidation are significantly enriched in AAV-Yap1 muscle (Fig. 5 E-H). Elevated BCAA catabolism, lipid transport and fatty acid oxidation suggest the generation and release of free BCAAs and oxidized lipids from the atrophying muscle. To verify this possibility, free BCAA and oxidized lipid levels were measured in blood plasma from Yap1^ON^ and Yap1^OFF^ mice. Yap1^ON^ mice showed a significant increase in blood plasma free BCAA and oxidized lipids compared to Yap1^OFF^ animals (Fig. 6I-J).

**Fig. 5:**
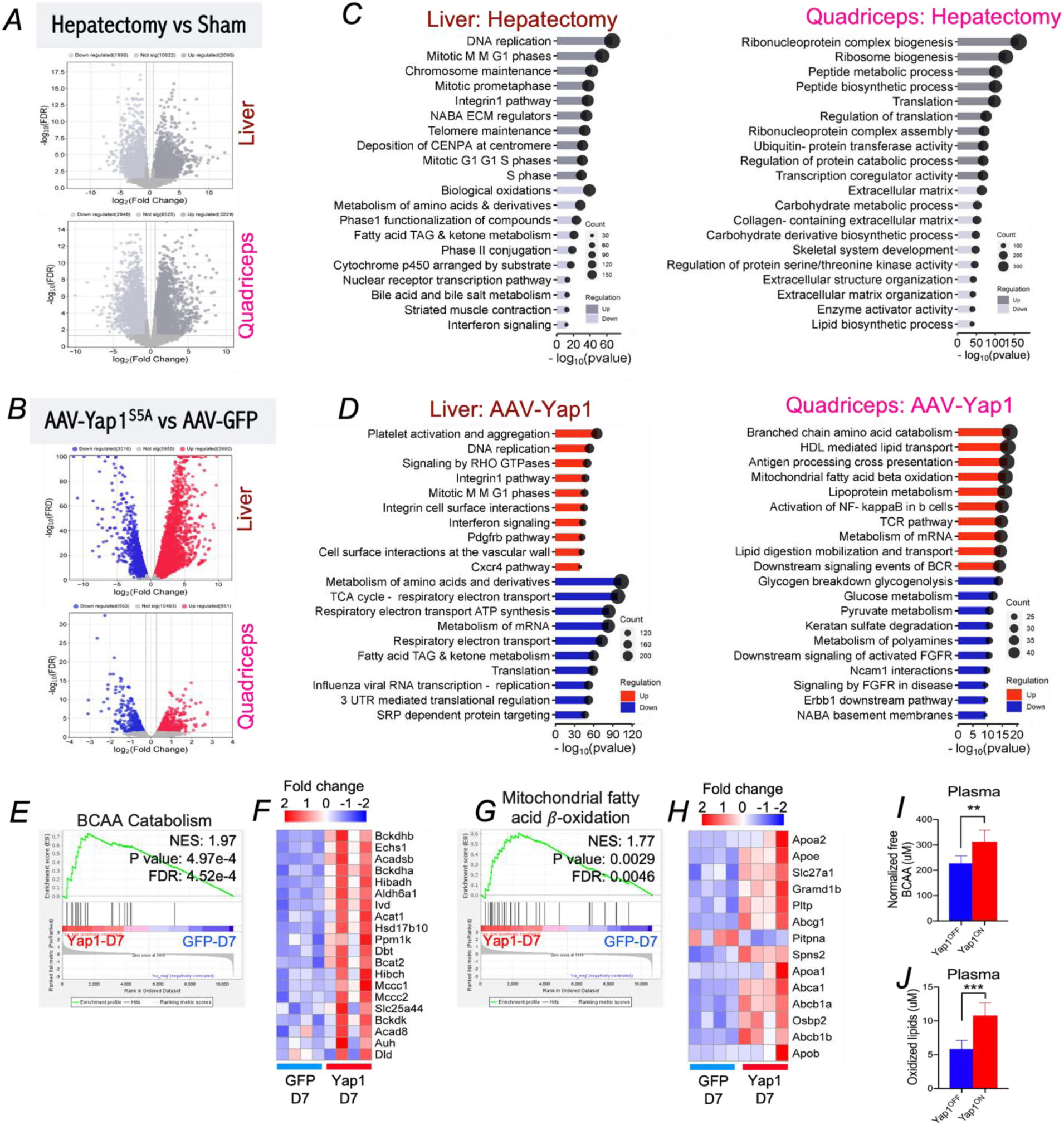
Liver and skeletal muscle gene expression in the liver growth mouse models. A) Volcano plot showing differentially expressed genes (DEGs) in whole livers and quadriceps from post-PHx mice. A cutoff of FC±1.5 and FDR≤0.05 was used. B) Volcano plot shows DEGs in whole livers and quadriceps from AAV-Yap1 injected mice. C) Top 10 canonical pathways significantly changed in livers and quadriceps after post-PHx, ranked by P-value. Dot sizes represent the gene number per canonical pathway. Left panel shows liver data, right panel shows quadriceps data. Upregulated and downregulated pathways are shown in light and dark gray respectively. D) Top 10 canonical pathways significantly changed in livers and quadriceps from AAV-Yap1 mice at day7 post-infection, ranked by P-value. Upregulated and downregulated pathways are shown in red and blue respectively. E) Gene set enrichment analysis (GSEA) of the BCAA catabolism in AAV-Yap1 quadriceps. F) Heatmap shows the DEGs within the BCAA catabolism signature in (E). Color scale shows fold change in gene expression. G) GSEA of mitochondrial fatty acid beta-oxidation in AAV-Yap1 quadriceps. H) Heatmap shows DEGs within the mitochondrial fatty acid beta-oxidation signature in (G). I) Free BCAA levels in blood plasma of day14 water fed, versus doxycycline fed *ApoE-Yap1* mice. n=5 mice per condition analyzed. **P<0.01, unpaired t-test. J) Oxidized lipids level in blood plasma of day14 water fed, versus doxycycline *fed ApoE-Yap1* mice. ***P<0.001, unpaired t-test.

**Fig. 6:**
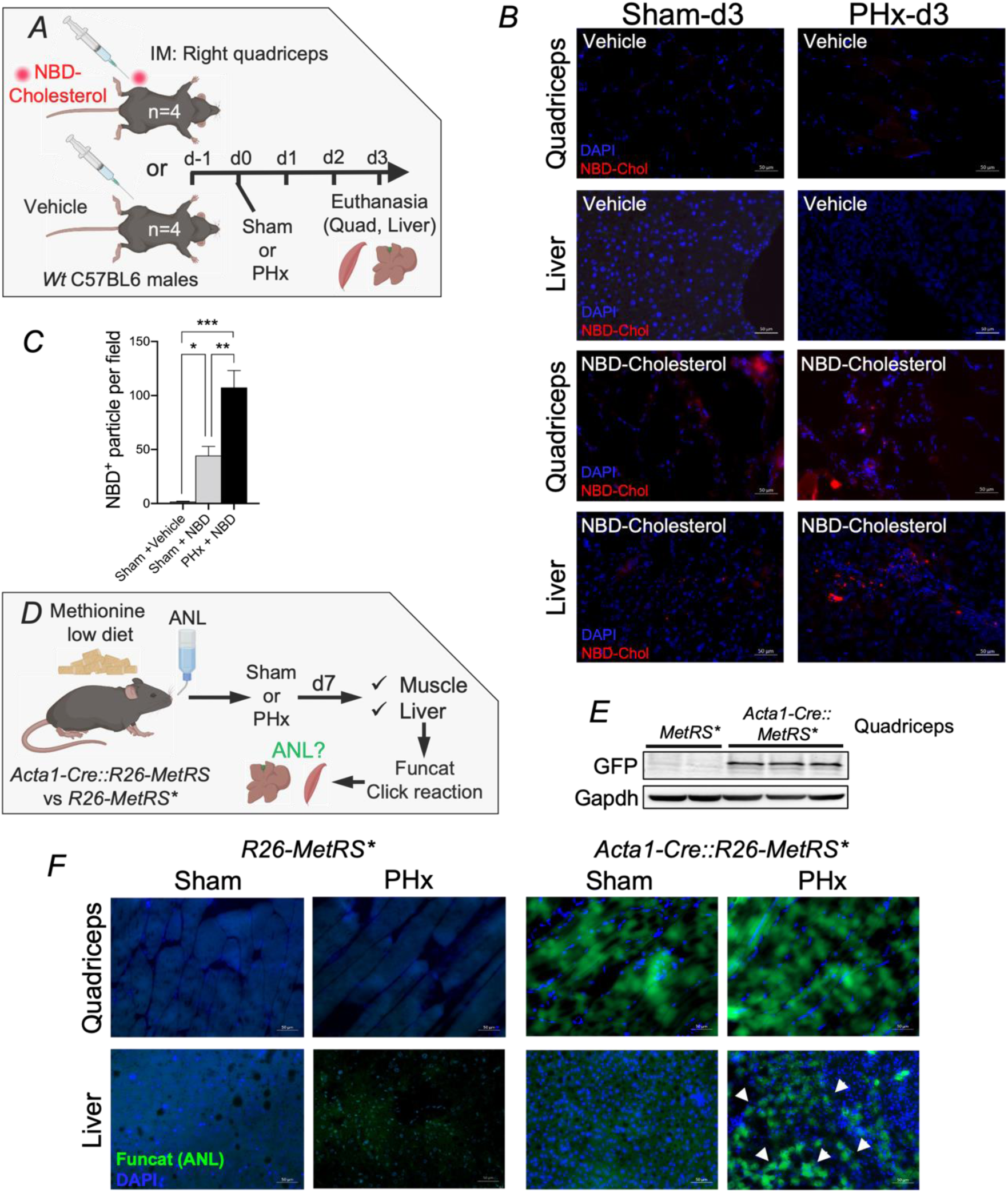
Muscle-derived cholesterol and amino acids relocate to the regenerating liver. A) NDB-Cholesterol tracing procedure. Mice (n=4 per condition) were injected with a 100ug of NBD-cholesterol or vehicle in the right quadriceps, 24hrs prior to surgery. The next day, sham surgery or 70% hepatectomy were performed in each group. Animals were euthanized on day3 post-surgery. B) Fluorescent microscopy detection of NBD-Chol on post-PHx livers and quadriceps described in (A). Cryo-sections were counterstained with DAPI to reveal nuclei. Scale bars= 50 microns. C) Quantification of NBD-Chol particles in the liver. Sham + vehicle, sham + NBD-chol and PHx + NBD-Chol livers were analyzed. *P<0.05, **P<0.01, \*\*\**P*<0.001, One-way ANOVA. D) ANL incorporation to skeletal muscle using the *Acta1-Cre::R26-MetRS\** mouse model. 3 weeks prior to surgery, animals were fed a methionine-low diet and given the methionine analog (ANL) for 10 days in drinking water prior to surgery. Sham and PHx surgeries were performed. Animals were euthanized at day7. Livers and quadriceps were collected and prepared for FUNCAT staining. E) Western blot analysis of GFP in *MetRS\** and *Acta1-Cre::MetRS\** quadriceps to validate the presence of GFP only in *Acta1-Cre::MetRS\** quadriceps. F) Representative images of FUNCAT staining (detection of ANL) on liver and quadriceps sections. White arrowheads show a strong hepatic ANL signal. Scale bars= 50 microns.

Similarly, free BCAA and oxidized lipids were found significantly elevated in the blood plasma, post-PHx (Supplementary Fig. 5). These findings suggest that muscle-derived BCAA and fatty acids may benefit liver growth.

### Muscle-derived amino acids and cholesterol relocate to the regenerating liver

Given the release of muscle-derived BCAAs and oxidized lipids in both liver growth models, we next traced muscle-derived lipids and amino acids in post-PHx livers. Fluorescent NBD-labelled cholesterol has been successfully used to trace lipids in the liver ^40^. NBD-Cholesterol was locally injected into the mouse quadriceps prior to PHx (Fig. 6A). At day 3 post-PHx, quadriceps and regenerating livers were collected, the NBD-Cholesterol content was measured. Quadriceps and livers from animals injected with vehicle did not show any measurable NBD-cholesterol, in either Sham or PHx mice (Fig. 6B). As expected, NBD-Cholesterol injected quadriceps showed accumulation of red fluorescent droplets in animals treated with either Sham or PHx. At day 3 post-surgery, NBD-Cholesterol was detected in sham non-resected healthy livers, confirming a well-established physiological exchange of fat between skeletal muscle and liver ^41,42^. Importantly, accumulation of NBD-Cholesterol droplets was 2-fold higher in post-PHx livers (Fig. 6B-C). These data demonstrate that muscle-derived cholesterol relocates at a higher extend to the regenerating livers. To trace muscle-derived amino acids in regenerating livers, the *STOPflox-R26-MetRS\** mouse model was used, as it allows tissue-selective labelling of nascent proteins with a methionine analog, azidonorleucine (ANL) ^43^. ANL can be detected in tissues using azide-alkyne click chemistry ^44^ (Fig. 6D). To initiate skeletal muscle specific translation of ANL labelled proteins, *Acta1-Cre::R26-MetRS\** mice were fed with a methionine-depleted diet and given ANL in drinking water prior to PHx. *R26-MetRS\** mice, (*Acta1-Cre^-^*), do not express mutant *MetRS* and therefore are not able to produce ANL labelled proteins, even in presence of ANL, and were used as a control group. Since the *MetRS\** mouse model carries a 2A-linkled GFP cassette, Acta1-Cre-mediated recombination of the floxed stop domain was verified in skeletal muscle by the presence of GFP in *Acta1-Cre::R26-MetRS\** quadriceps extracts but not in *R26-MetRS\** (Fig. 6E). Seven days after surgery, fluorescent non canonical amino acid tagging (FUNCAT) was performed on muscle and liver sections as previously described^44^. FUNCAT signal was detected in *Acta1-Cre::R26-MetRS\** quadriceps in both sham and PHx-mice clearly showing incorporation of the amino acid analog ANL into skeletal muscle, whereas no signal was detected in *R26-MetRS\** quadriceps (Fig. 6F). FUNCAT signal was detected only in post-PHx livers from *Acta1-Cre: R26-MetRS*\* whereas sham livers from both genotypes were negative for FUNCAT signal (Fig. 6F). Collectively, our findings demonstrate that muscle-derived amino acids and lipids relocate to the regenerating livers in vivo.

### Hepatic and muscle gene expression in liver growth models is highly similar to cancer models

We showed that changes in liver homeostasis, as a consequence of partial hepatectomy or Yap1-induced hepatomegaly, induce a cachexia phenotype, similar to what is observed in various cancer cachexia mouse models. For instance, pancreatic ductal adenocarcinomas, Colon 26, Lewis lung carcinomas and liver metastasis mouse models all show severe body weight loss, muscle atrophy and fat loss and constitute suitable models to investigate cancer cachexia ^45–49^. To assess the genetic similarity between liver growth models and various cancer models, we performed a correlation analysis of gene expression changes in livers and muscles. Livers from PHx and AAV-Yap1 models showed the most significant gene expression correlation with liver tissues from pancreatic cancer (orthotopic KPC and KIC), melanomas bearing mice, liver metastasis, and c-Met-induced hepatocellular carcinomas mouse models, with the highest correlation score when compared to livers from orthotopic KPC (Fig. 7A, C). Similarly, muscles from PHx and AAV-Yap1 models showed highly similar gene expression patterns with muscle from pancreatic cancer (orthotopic KPC and genetic KPC), metastatic colon cancer, C26 colon cancer and Lewis lung cancer models, with the highest correlation score against orthotopic KPC muscle (Fig. 7B-D). The full correlation analysis data is available in the Supplementary table 2. Based on the top ranked gene expression correlations in (Fig. 7A-D), we chose to further compare gene expression patterns in PHx and orthotopic KPC. Top 200 marker genes, selected based on a signal-to-noise criteria, from PHx and orthotopic KPC tissues were clustered into heatmaps (Fig. 7E, F). Hepatic gene expression heatmap confirms that gene expression changes in post-PHx livers highly resemble those observed in livers from mice bearing a pancreatic tumor (Fig. 7E). Moreover, the top 5 commonly affected signaling pathways in PHx and KPC livers were related to the extracellular matrix, metabolism, complement, cytokines and Il6/Jak/Stat3 signaling, highlighting the similar changes in liver homeostasis and metabolism after injury (PHx), as well as in presence of a pancreatic tumor. Likewise, muscle gene expression heatmap confirms greatly similar gene expression patterns in post-PHx and pancreatic tumor bearing mice (Fig. 7F). Commonly affected genes were related to upregulated functions such as proteasome degradation, protein catabolic process and downregulated functions, such as myogenesis and extracellular matrix, generally associated with muscle atrophy in many other contexts ^50^. Altogether, our findings demonstrate the existence of a hepatic and muscle genetic response to the presence of liver growth and reparative signals, similar to those observed in the presence of tumor legions. While a similar gene signature in the muscle is predictable, due the predominant atrophy phenotype, in the liver however, our data suggest that solid tumors remotely modify the liver genetic program in a similar fashion observed during liver growth.

**Fig. 7:**
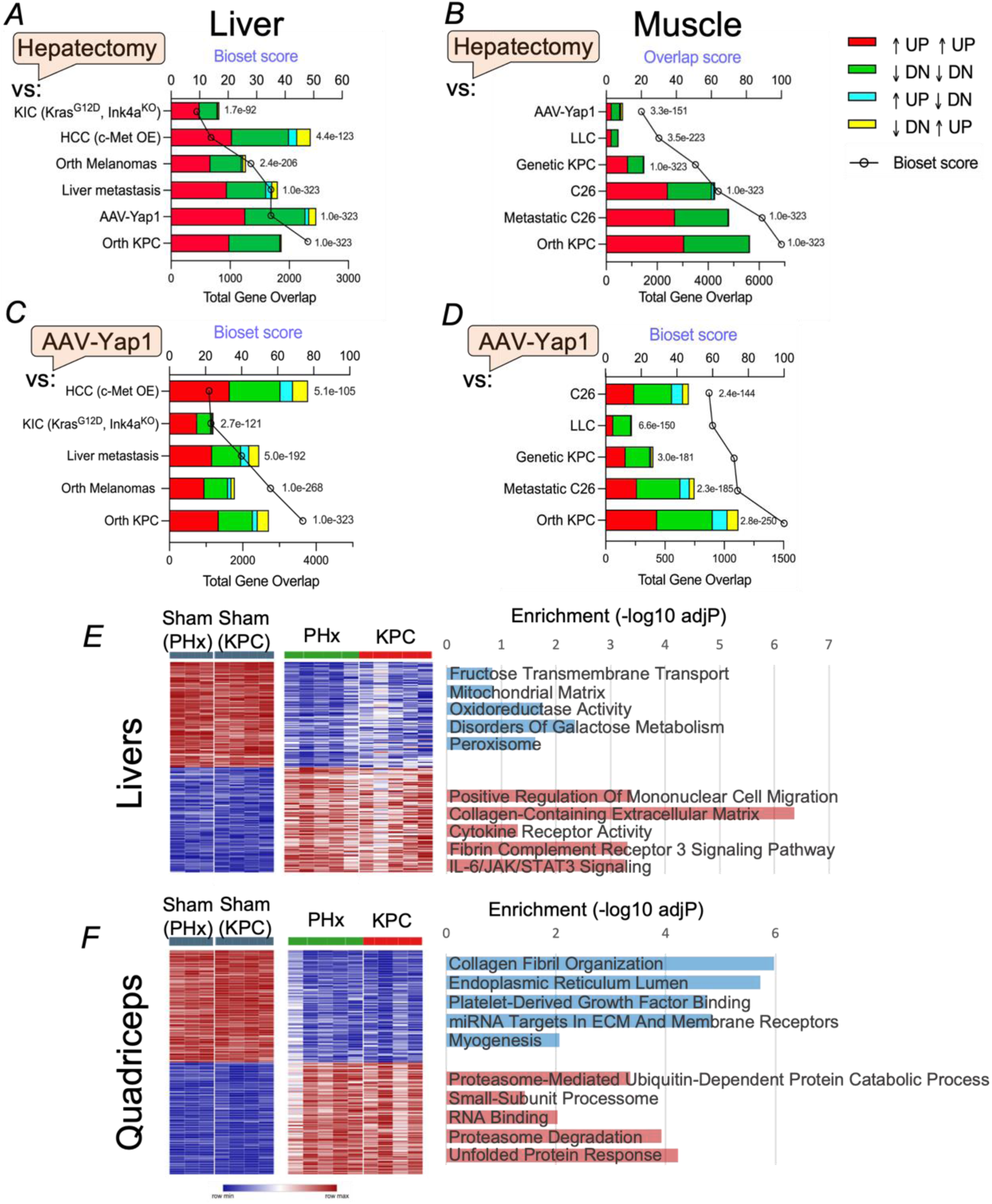
Hepatic and muscle gene expression in liver growth models is highly similar to cancer models. A) Correlation analysis of gene expression in livers from post-PHx mice, versus livers from cancer mouse models that induce muscle atrophy. The number of commonly upregulated and downregulated genes are plotted in red and green histograms respectively. Genes expressed in different directions (Up in PHx ˂-˃ Down in others, or Down in PHx ˂-˃ Up in others) are plotted in blue and yellow histograms. Correlations are ranked by overlap score and overlap p-Value. B) Correlation analysis using DEGs in post-PHx muscles. C) Correlation analysis using DEGs in AAV-Yap1 livers. D) Correlation analysis using DEGs in AAV-Yap1 muscles. E,F) Gene expression heatmap of liver expression patterns for 200 marker genes (100 positive and 100 negative) selected from the transcriptomic analysis, distinguishing livers from PHx model and orthotopic KPC model and their respective controls. (E) shows liver transcriptomic data and (F) shows muscle data. Marker gene selection was based on signal-to-noise criteria using the broad institute Morpheus webtool. Right panel, bar graphs represent the top 5 enriched terms from GO Biological Process, GO Cellular Component, GO Molecular Function, WikiPathway 2023, and MSigDB Hallmark 2020, ranked by the most significant adjusted p-value. Blue bars indicate decreased terms in both PHx and KPC compared with Sham, while red bars represent increased terms in both PHx and KPC compared with Sham controls.

## Discussion

This study examines the regulation of muscle mass during the hepatic proliferative response. We identified the hepatic metabolic regulator Yap1 as a systemic mediator of both muscle loss and catabolic state during liver regeneration. Liver regenerative response results in an approximate 25 % loss of muscle mass and over a 50 % loss of fat mass with increased REE and utilization of a mixed oxidative fuel source. In vitro studies demonstrate that the loss of muscle mass can be recapitulated in fused myotubes treated with plasma, or conditioned media from regenerating liver, and in vivo with the hepatocyte specific expression of a constitutively active Yap1. Thus, hepatic Yap1 integrates liver regeneration to the secretion of serum factors that directly induce muscle loss. In vivo tracer studies further demonstrate that hepatic regeneration occurs with the transfer of substrates including amino acids and lipids from muscle to liver. Thus, muscle loss likely occurs to provide building block substrates to the regenerative response. Consistently, following a 45 % resection of lung mass, a major surgical trauma without a regenerative response, no loss of fat or muscle was observed. We unraveled a new role for Yap1 transcription factor in promoting liver repair and size control, via the induction of a cachexia phenotype in mice. In this physiological cachexia mechanism, Yap1 promotes the release of liver factors to induce muscle atrophy and fat loss in mice (Fig. 8). Comparing liver transcriptomes in liver growth and pancreatic cancer model, we found great similarities in their respective hepatic gene expression patterns. Therefore, we believe that the presence of cancer lesions in mice triggers similar changes to the liver homeostasis and metabolism, thus, diverting the reparative physiological cachexia to sustain tumorigenesis.

**Fig. 8:**
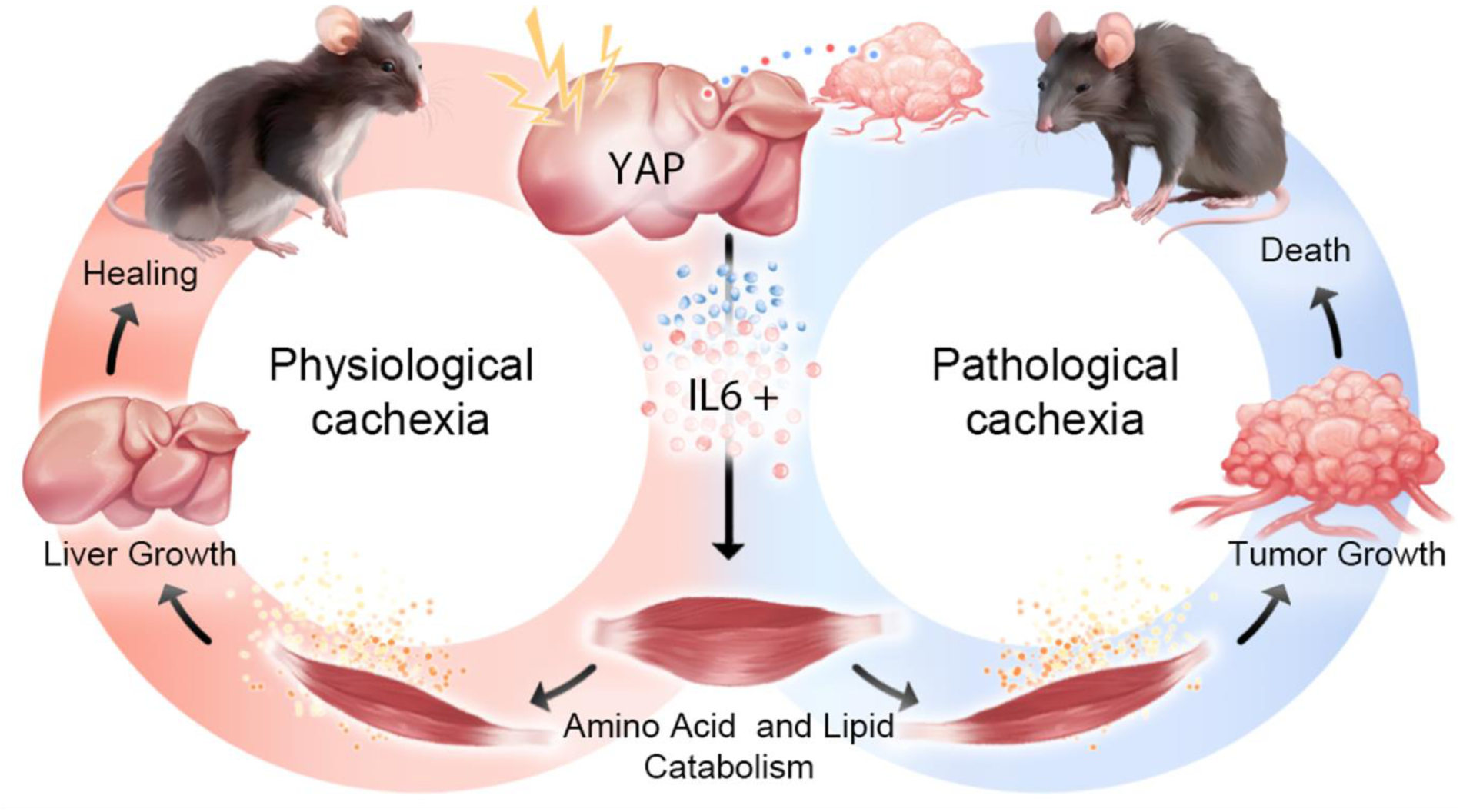
Model of beneficial versus detrimental effects of cachexia. Liver reparative and growth processes induce a physiological cachexia in mice, including a rapid body weight loss, muscle and fat loss. Physiological cachexia is transient, reversible and promotes liver repair and growth. the transcriptional co-activator Yap1 is involved in physiological cachexia by inducing a systemic catabolism and the release of hepatic secreted factors that trigger muscle atrophy. Muscle atrophy in turn, releases metabolites (Amino acids and lipids) that fuel liver repair and growth. In presence of pancreatic tumors, a tumor/liver crosstalk is established, and induces changes in liver genetic expression, similar to those observed during liver repair and growth. Our findings suggest that tumor growth utilizes the liver-induced physiological cachexia to support the tumor growth. In contrast to physiological cachexia, this tumor-driven pathological cachexia is non-reparative and detrimental to animals health.

It is well documented since the 1930s that the healing process that accompanies a major trauma, surgery, or other reparative processes requires substrates to support ongoing metabolic needs but to also rebuild and repair tissue. Studies over many decades since this initial report have observed negative nitrogen balance following various injuries ^51,52^. Further nutritional studies have identified that the reparative process generally is not directly supported by enteral nutrition but rather requires the body to activate a catabolic state that results in skeletal muscle and fat loss. Further, many of these injury responses across species are associated with a temporary ileus and thus the inability to nutritionally meet metabolic needs or even tolerate food.

Here, hepatic Yap1 and Taz are found to integrate the regenerative liver response to the induction of muscle and fat catabolism. Mice that carry *Yap1/Taz* deficient livers failed to restore liver mass after partial hepatectomy, due to a defective hepatocyte proliferation^15^. Pan and coworkers similarly found that overexpression of Yap1 was associated with massive hepatomegaly ^21^. The studies reported herein, extend these observations and demonstrate that liver expansion occurs in association with loss of muscle and fat mass. Furthermore, we demonstrate that hepatic Yap1 activation is sufficient to change the metabolic state of the mouse. Yap1 hepatic activation increased REE and decreased the RQ.

Liver growth models caused gene expression changes in skeletal muscle and livers highly similar to those triggered by several cancer cachexia models. Hepatic alterations associated with many types of cancer are thought to be the main cause of metabolic changes leading to cachexia ^53,54^. Moreover, chronic liver disease and hepatocellular carcinomas (HCC) also correlate with a severe loss of muscle mass and function ^55,56^. Although similar gene expression changes in the skeletal muscle are expected from mouse models all inducing muscle atrophy, gene expression similarities between regenerative livers and livers from cancer cachexia models raise the possibility that cancer may commandeer liver homeostasis to cause cachexia and pathologically grow from the nutrients released by muscle and fat wasting.

Yap1 is a prognostic marker in chronic liver disease and HCC ^57–59^. Our study therefore also suggests that Yap1 may govern the shift to a catabolic state in cirrhotic patients. Interestingly, blood plasma from our Yap1 infected animals induced a significant myotube atrophy in vitro. This suggests that Yap1 drives a secretory program in the liver that promotes muscle atrophy. We tested two secreted candidates (*Il-6* and *Ctgf*), known as Yap1 target genes, as potential mediators of muscle and fat loss with negative results. Proteomic characterization of the Yap1-mediated secretome may help identify new muscle and fat loss mediators that may be therapeutically exploited to target body weight loss in various pathologies.

To date, most nutritional strategies for cancer and liver disease patients have had only a minimal impact on cachexia ^8,60,61^. Our study shows that skeletal muscle-derived metabolites relocate to the regenerative liver during the repair in a process that, based on the muscle gene expression pattern, is greatly similar from muscle in cancer models. As the main metabolic center, liver operates a constant physiological crosstalk with many organs to control lipid, protein and glucose metabolism ^62,63^. We believe that, during liver repair, a fine-tuned liver-muscle crosstalk is established and promotes liver regeneration via the release of nutrients, energy, and metabolites, at least in part from muscle breakdown. This reparative liver-muscle crosstalk appears to be a conserved mechanism predominant to the nutritional support and may represent a better target to overcome unvoluntary weight loss in many pathologies.

## Methods

### Animal models

Animal experiments were approved by the Indiana University, and Oregon Health and Science University Institutional Animal Care and Use Committees (IACUC). Mice were housed in a barrier facility with ad libitum access to autoclaved food and sterile water, maintained on a constant temperature on a 12-h light/dark cycle. All animals used for this study were male mice aged between 12 and 14 weeks old. C57BL6 mice were allowed to acclimate to the facility for, at least 1 week before use. To induce hepatic Yap1^S5A^ expression, *ApoE-rtTA-Yap1^S5A^*mice were given doxycycline 0.2mg/ml (Sigma-Aldrich) in drinking water. To generate the skeletal muscle specific MetRS* line, the *STOPflox R26-MetRS\** mice (Jackson laboratory, #028071) were crossed to the *Acta1-Cre* mice (Jackson laboratory, #006149). Animals homozygous for MetRS* were used. Animals were fed with a methionine-depleted diet ad libitum (ResearchDiets, A05080226i) for 4 weeks and given L-azidonorleucine (Tocris, #6585) 60uM in drinking water for 2 weeks prior to the experiment.

### Partial hepatectomy

70% hepatectomy (PHx) was performed as previously described [16]. For analgesia, mice were administered Ethiqua XR, 3.25mg/kg the day of surgery. Mice shaved and anesthetized with isoflurane 2-4% and operated under a 37°C heat pad with eye ointment. Betaine was applied 3 times on the shaved surgical area. An upper sternal laparotomy was performed. 4-0 nylon suture (Ethicon, #662G) was used to tie and resect the left lateral and medial lobes. Laparotomy was sutured with isopropanol soaked 4-0 silk surgical sutures (Ethicon, #781B). Skin was closed with 9mm wound clips (Alimed, # MPSAM-98SUT25-6). Mice were allowed to recover on 34C heat pads (Braintree Scientific, TP-700) for 24 hours. Disease score was monitored daily for up to 4 days after surgery.

### Adeno-associated virus (AAV)

AAVs (Vector Biolabs, PA) were dissolved in sterile PBS according to the manufacturer’s suggested dilutions. *AAV8-TBG-eGFP* and *AAV-DJ8-TBG-Yap1^5SA^* were injected under anesthesia retro orbitally at 2.52×10^12^ GC. *AAV-DJ8-TBG-mCTGF* were delivered at 1.2×10^12^ GC. *AAV-Null* (empty) and *AAV-CAG-mIl-6* were injected at 1.2×10^12^ GC.

### Euthanasia and tissue collection

Animals were anesthetized with isoflurane 2-4%, then cervically dislocated. For blood collection, animals were anesthetized and a cardiac puncture was performed to collect blood in an EDTA vacutainer® (BD). Animals were then cervically dislocated. Organs were collected, weighted and stored accordingly for subsequent applications. Tissues were either flash frozen in liquid nitrogen and cryopreserved at-80°C, fixed in formalin 10%, or embedded in OCT after isopentane snap freeze in dry ice.

### Muscle fiber CSA and frequency distribution measurement

Muscle fibers were first stained with dystrophin. CSA was measured from 20 random fields using an ImageJ macro that automatically contrasts the dystrophin ^64^. For each cross section, 300-400 myofibers were measured, and a mean was calculated.

To measure the frequency distribution of CSA area, 1800 to 2400 fibers were measured in each condition. Prism software was used to calculate and bin the frequencies in each condition.

### Grip test and hanging

Grip strength test was performed using the commercially available automatic grip strength meter (Bioseb grip strength meter). The four-limb hanging test was conducted using a wire grid system to noninvasively measure the ability of mice to exhibit sustained limb tension to oppose their weight.

### Liver lobe media conditioning

Media conditioning with post-PHx livers were performed as previously described ^65^. Post-PHx or sham liver lobes were weighted, washed once with ice cold PBS, incubated in ice cold Krebs buffer, and transferred to a 6-well plate well filled with 3ml of myotube differentiation media. The next day, conditioned media (CM) was collected, 0.2u filtered and added to C2C12 myotubes. The volume of CM added to myotubes was proportional to the initial weight of each liver love used.

### Muscle atrophy in vitro assay

C2C12 mouse myoblasts were a curtesy of Dr. Paola Costelli (University of Turin, Turin, Italy). Myoblasts were maintained in DMEM (Corning, 10-017-CV), 10 FBS (Cytiva, SH30396.02) supplemented with penicillin/streptomycin cocktail (Cytiva, SV30010).

For differentiation into myotubes, C2C12 myoblasts were seeded at around 90% confluence on day1. On day2, Cells were washed once with warm PBS and growth media was replaced by the differentiation media, DMEM, 2% horse serum (Cytiva, SH30074.04HI) supplemented with penicillin/streptomycin. Media was replaced every other day. 5 days differentiated myotubes were used for experiments.

### Western blotting

Standard SDS-PAGE was used to detect the candidate protein. 4-15% gradient acrylamide gels were used to run 30ug of total proteins. Proteins were transferred into a nitrocellulose membrane. IRDye secondary antibodies (LI-COR) were used. Immunoblot was revealed using the Odyssey CLx (LiCor). A full list of antibodies used is available in the supplementary table 3 (Antibody_list).

### Real-time quantitative PCR

Total RNAs were extracted with the QIAzol lysis reagent (Qiagen, 79306) and cleaned up with the RNA clean up kit (Qiagen, 74204). cDNA was generated using the iScript™ cDNA Synthesis Kit (Biorad, 1708890). Light Cycler 96 (Roche) was used to quantify the cycle threshold (Ct). A list of the primers used is available in the supplementary table 4 (Primers_list).

### Immunohistochemistry and immunofluorescence

Immunohistochemistry was performed as previously described ^66^. 6µm FFPE sections were used. Antigen retrieval was performed in a 96°C water bath using EDTA, pH9 buffer. ImmPACT DAB kit from Vector Laboratories (SK-4105) was used for detection. Sections were counterstained with hematoxylin prior to dehydration and mounting in Cytoseal® (ThermoFisher, 831016). For cryosections, standard IF was used. Cryosections were fixed in 4% paraformaldehyde, permeabilized in 0.01% triton X100, blocked in 10% goat serum (Sigma, G9023). Primary antibodies were applied overnight in cold room. Secondary antibodies were applied for an hour at room temperature. Prolong gold DAPI (Invitrogen, P36935) was used to mount slides. For C2C12 myotubes, IF was performed in a 12 well plate. Counterstaining was done with NucBlue™ nuclei stain (Invitrogen, R37605). Images were obtained using a Zeiss epi-fluorescent microscope and Lionheart LX automated microscope (BioTek).

### RNA-Sequencing, GSEA and correlation analysis

Based on the significance of liver mass and muscle mass change post AAV-Yap1 infection, gene expression in both livers and quadriceps was analyzed at day7 post AAV infection. Post-PHx tissues were analyzed at day1, day3 and day7 post-surgery. Based on the earliest detection of muscle atrophy, and total DEGs in this model, we chose to analyze tissue gene expression from day3 post-PHx mice. Single-indexed strand-specific cDNA library we obtained using TruSeq Nano DNA Library Prep kit. Library quantity and size distribution using a Qubit and Agilent 2100 Bioanalyzer. 200 pM pooled RNA libraries were used per flow cell for clustering amplification on cBot using HiSeq 3000/4000 PE Cluster Kit and sequenced with 2 × 75–bp paired-end configuration on a HiSeq4000 (Illumina) using the HiSeq 3000/4000 PE SBS Kit. Q score was used to measure sequencing quality. More than 95% of the sequencing reads reached Q30 (99.9% base call accuracy). Sequencing data were first assessed using FastQC (Babraham Bioinformatics). Sequenced libraries were mapped to the mm10 mouse genome using STAR RNA-seq aligner ^67^. Uniquely mapped sequencing reads were assigned to the mm10 UCSC reference genome. Data were normalized using the trimmed mean of M values method. Differential expression analysis was performed using edgeR and the FDR was computed from P values using the Benjamini-Hochberg procedure. The chosen cuttoff was (FC of ≥1.5, FDR of ≤0.05). Data was deposited at the gene expression omnibus (GEO) under accession number GSE262617 and GSE262620 for AAV-Yap1 and PHx respectively. BaseSpace Illumina was used for gene ontology and canonical pathways analysis (Supplementary table 1 and 2). GSEA was used to further confirm the enrichment of selected pathways. Correlation analysis was performed using BaseSpace Illumina application. Data was uploaded to Illumina BaseSpace Correlation Engine 2.0. A Meta-Analysis comparing livers and muscles from the liver growth models to publicly available biosets was performed. Bioset scores were calculated by overall statistical significance and consistency of the enrichment, or overlap, between the genes or SNPs in our biosets (liver growth) and queried biosets. Genes upregulated in both biosets, downregulated in both biosets, and conversely regulated between biosets contribute to the total gene overlap. P-value is displayed to the right of the total gene overlap. Public datasets of muscle atrophy models identified by the correlation analysis are listed below.

### Public datasets found by correlation analysis

Liver and muscle expression data from our liver growth models were most correlated with the following studies: Genetic KPC: GSE157251. Metastatic C26: GSE142455. C26: GSE142455. Orthotopic KPC: GSE123310 and GSE157251. Orthotopic melanomas: GSE199863. KIC: GSE51931. HCC: GSE25142.

### Web tools

SRplot webtool ^68^ was used to generate volcano plots, canonical pathways bar graphs, selected pathway heatmaps and comparative gene ontology bar graphs. Morpheus, https://software.broadinstitute.org/morpheus, was used to perform top 200 marker gene selection based on signal-noise ratio and heatmap generation while comparing PHx and KPC datasets. Enrichr ^69^ was used to generate the top 5 enriched terms from GO Biological Process, GO Cellular Component, GO Molecular Function, WikiPathway 2023, and MSigDB Hallmark 2020 while comparing PHx and KPC datasets.

### Fluorescent non-canonical amino acid tagging (Funcat)

Funcat was performed as previously described ^70^. 6µm cryosections were fixed in in 4% paraformaldehyde, permeabilized in triton X100, blocked with 1% FBS for an hour at room temperature. Click chemistry reagents were mixed in PBS with intermittent vortex as follows: THPTA (Sigma-Aldrich, 762342, 3mM), Alexa Fluor 488 Alkyne (Invitrogen, A10267, 2uM), sodium ascorbate (Sigma-aldrich, 1613509, 5mM), CuS04 (Sigma-Aldrich, 451657, 600 uM). Slides were incubated with the click mixture in the dark for 4 h, then washed twice with PBS-Tween 0.5 mM EDTA solution for ∼ 15 min. One last wash with plain PBS was applied. Prolong™ gold with DAPI was used to counterstain nuclei and mount the slides.

### NBD-cholesterol tracing

Mice were anesthetized and the skin was shaved and wiped with 70% ethanol. 100ug of NBD-Cholesterol (ThermoFisher, N1148) dissolved in 100% ethanol (Sigma, #E7023) was injected using a 23G needle into the right quadriceps muscle in two distant spots (2×50ul) to avoid leaking of material. Vehicle animals were injected 2×50ul of 100% ethanol. Mice recovered on a heat pad and were used for partial hepatectomy, 24 hours after NBD-Cholesterol or vehicle injection.

### BCAA and oxidized lipids assay

Blood plasma BCAAs and oxidized lipids were quantified using the branched chain amino acids kit (VWR, 103721-714) and the TBARS, TCA method kit (Cayman chemical, 700870) respectively, following manufacturer’s instructions. Plasmas from 5 mice per condition were tested.

### Statistical analysis and data presentation

Data are presented as mean ± standard deviation (SD), unless otherwise specified. Tissue mass data is presented as percentage of initial body weight (%IBW). Cell culture experiments were performed at least two times, with three biological replicates and at least, two technical replicates each time. 70% hepatectomy was performed at least three time. Overexpression of Yap1^S5A^ using the *ApoE-rtTA-Yap1^S5A^*were replicated at least three times. AAV experiments were carried out at least two times. Western blot band intensities were quantified using ImageJ software and normalized to loading control band intensities. For *in vivo* studies, n=5 per cohort was chosen as an optimal number to appreciate a 10% statistical difference between conditions. Statistical analyses were performed using GraphPad/Prism 9.0. Sample size is specified in the figure legends. To test two datasets, unpaired two-tailed Student’s *t* test was used. To test the means among three or more datasets, one way, or two-way ANOVA with the Tukey’s multiple comparison test were used. The differences between groups were considered statistically significant when *P*<0.05. Significance code: \**P*<0.05, \*\**P*<0.01, \*\*\**P*<0.001, \*\*\*\**P*<0.0001, ns: not significant.

## Acknowledgement

This work was funded by the National Institute of Health: National Institute of General Medical Sciences, Grant/Award number: R01GM137656/GM/NIGMS.

Sequencing analysis was carried out in the Center for Medical Genomics at Indiana University School of Medicine, which is partially supported by the Indiana University Grand Challenges Precision Health Initiative. Metabolic cages experiment was carried out by Natalie Stull at the center of diabetes and metabolic diseases at Indiana University School of Medicine. Experimental design figures were created with BioRender.com. Working model (Figure 8) was created with the artistic help of Tetiana Korzun.

## Authors contribution to Manuscript

L.G.K. and T.A.Z. initiated the research project; L.G.K., T.A.Z. and C.D.W. assisted with study design and data interpretation; T.H. Y.J., T.R., A.K., E.A., J.E.R., X.Z., S.M., T.L. conducted experiments, acquired, analyzed and interpreted the data; A.R. and A.H. provided the MetRS* mice and assisted with the MetRS/FUNCAT experiments and guidance with data interpretation; R.F.C. and S.S.C. assisted with bioinformatics analysis; T.H. and L.G.K. wrote the initial draft manuscript and subsequent edits; C.D.W., J.E.R., A.R. and A.K. helped review and edit the manuscript.

## Data availability

RNA-Sequencing on AAV-Yap1 livers and muscles have been deposited to the gene expression omnibus database. Accession number: GSE262617. Reviewer token can be found in the link below: https://urldefense.com/v3/ https://www.ncbi.nlm.nih.gov/geo/query/acc.cgi?acc=GSE262617;!!Mi0JBg!J75CaHjlj1w8gyFzQrXNKZn5zyaqJsuXdzUDLk2yH7Xtdz2XsMo4J8kA2tDIK5RBezmjAqm5scf4-42s4Q$

Enter token ojubasoklbahnud into the box.

RNA-Sequencing on post-PHx livers and muscles (Day3) have been deposited to the gene expression omnibus database. Accession number: GSE262620. Reviewer token can be found in the link below: https://urldefense.com/v3/ https://www.ncbi.nlm.nih.gov/geo/query/acc.cgi?acc=GSE262620;!!Mi0JBg!OWCHfUUVZ34uyLjZLRdJHQ_9eZHzpMY44V31uPy9TMrJmlu61H5ez3QPgw3bGpF2cFWUMakX77dv5AQPxg$ Enter token yfgfyqswvdctvef into the box

## Material and correspondence

Dr Leonidas Koniaris and Dr Teresa Zimmers are the two authors to correspond with and contact for any material request.

---

**Supplementary Fig. 1:**
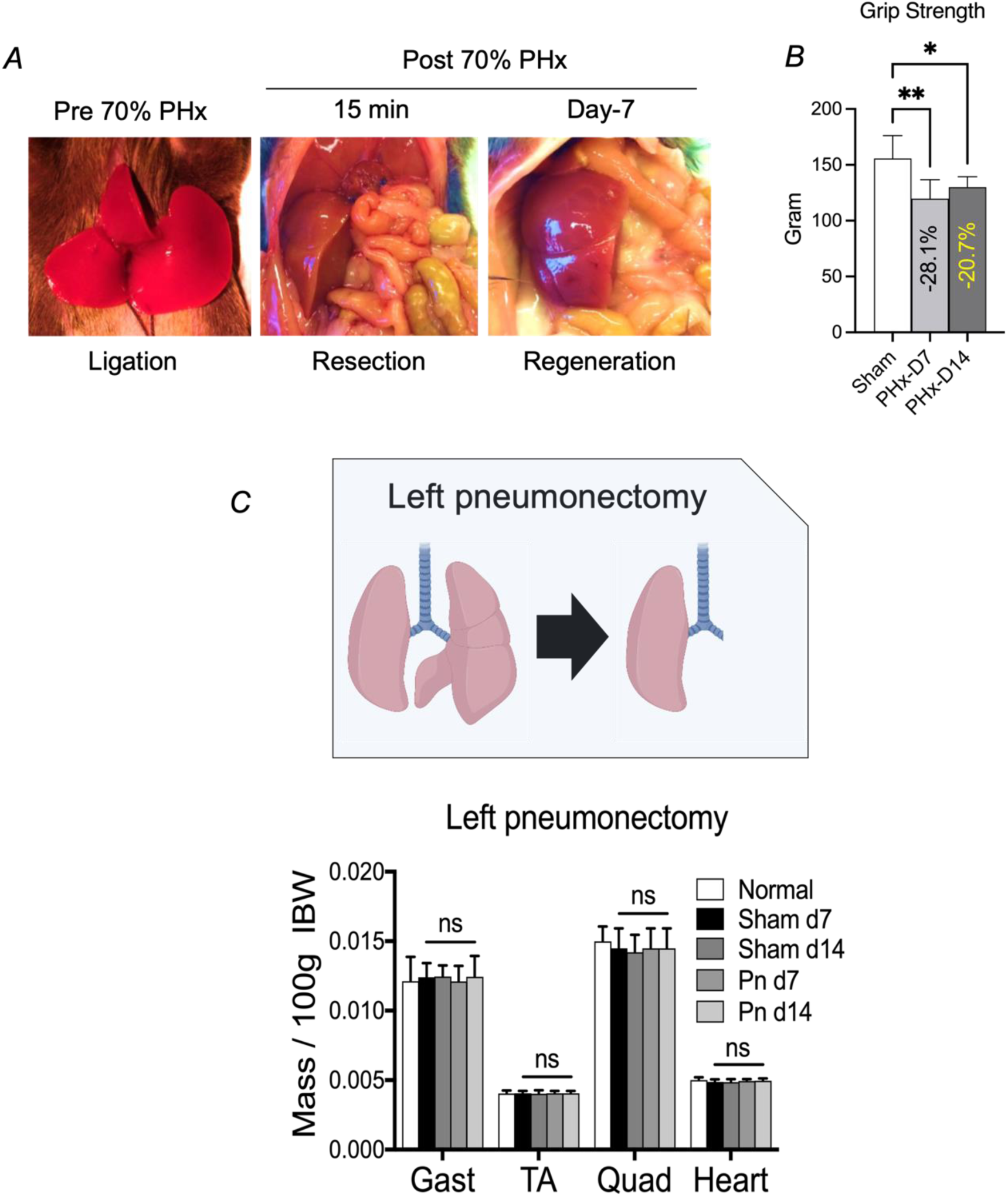
Regenerative (partial hepatectomy), versus non regenerative (Left pneumonectomy) surgery. A) Images of 70% partial hepatectomy (PHx) showing liver lobes ligation, resection and Day-7 liver regeneration. B) Grip test in day7 and day14 post-PHx mice. n=5 per condition, *P<0.05, **P<0.01, one-way ANOVA. C) Schematic representation of left pneumonectomy in mice. D) Skeletal muscle and heart weight at day7, day14 post-left pneumonectomy in mice. n=5 mice per condition, One-way ANOVA.

**Supplementary Fig. 2:**
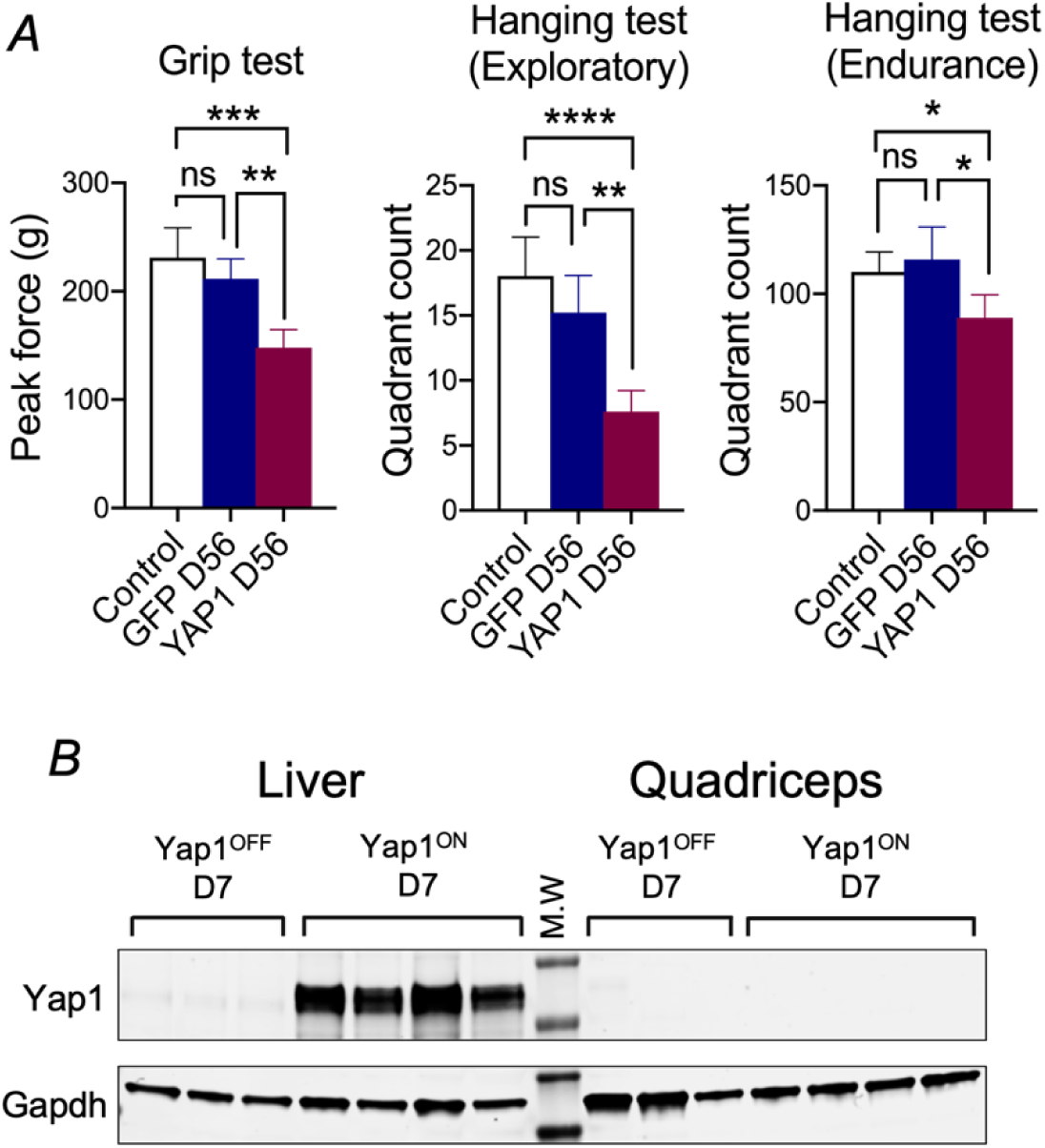
Yap1-induced liver growth causes muscle weakness. A) Grip test and hanging test in Control (No AAV injected), AAV-GFP and AAV-Yap1 injected mice, 8 weeks post-injection. n=5 mice per group. *P<0.05, **P<0.01, ****P<0.0001. B) Western blot analysis for Yap1 in liver and quadriceps extracts of water fed (Yap1^OFF^), versus doxycycline fed (Yap1^ON^) mice.

**Supplementary Fig. 3:**
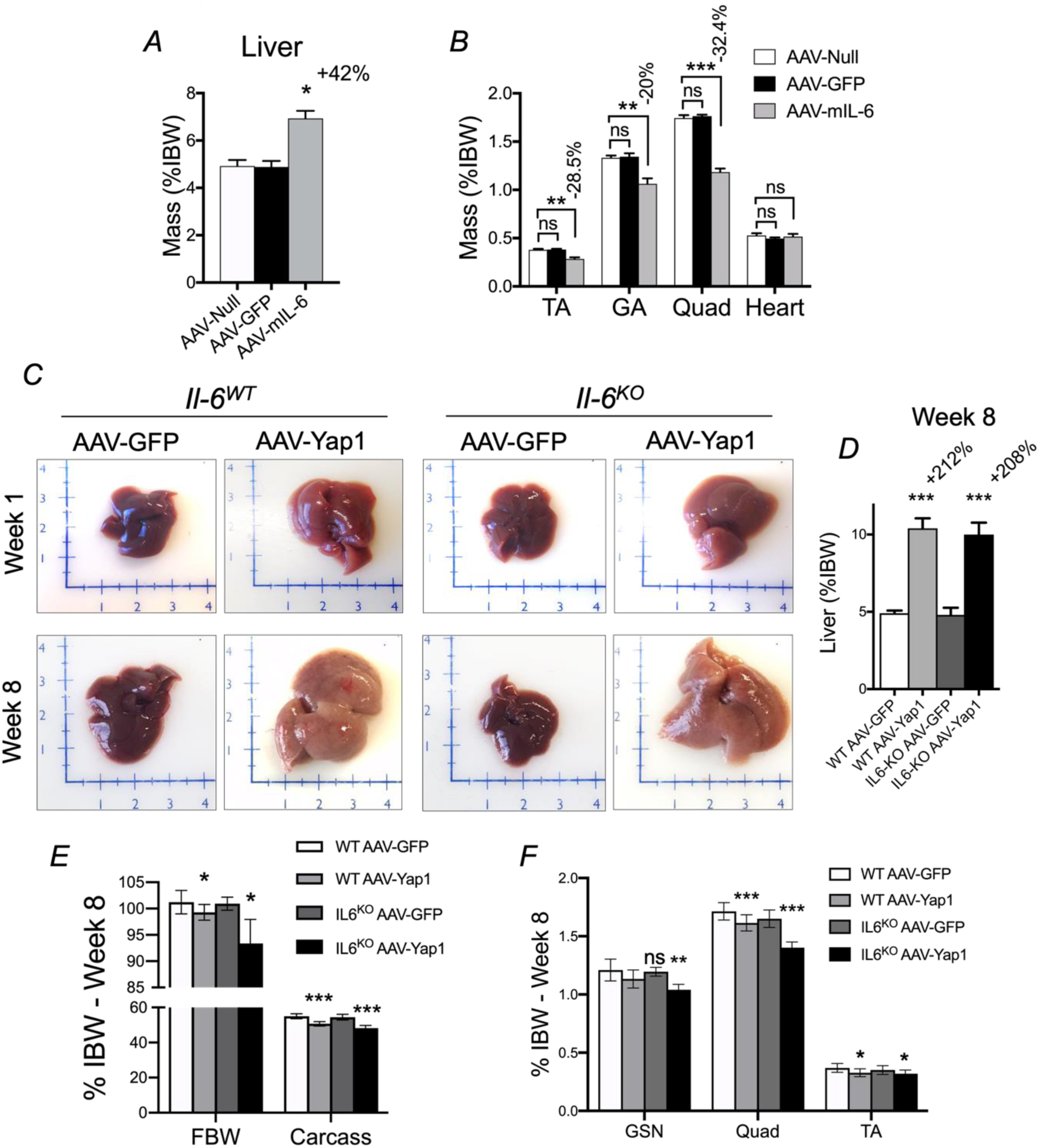
Yap1-mediated hepatomegaly induces body weight loss and muscle loss independently of Il-6. A) Liver mass in AAV-Null, AAV-GFP and AAV-CAG-mIl-6 infected mice. n=8 mice per condition, \**P*<0.05, one-way ANOVA. B) Muscle and heart mass of animals from (A). \*\**P*<0.01, \*\*\**P*<0.001, one-way ANOVA. C) Representative images of livers from AAV-Yap1 infected mice in both *Il-6^WT^* and *Il-6^KO^* genotypes, 1 week and 8 weeks post-infection. D) Liver mass in (C), presented as percentage of initial body weight (%IBW). n=5 per condition, \*\*\**P*<0.001, one-way ANOVA. E) Final body weight and carcass weight in each condition at week 8 post-infection. \**P*<0.05, \*\*\**P*<0.001, one-way ANOVA. F) Skeletal muscle mass at week 8 post-infection. \**P*<0.05, \*\**P*<0.01, \*\*\**P*<0.001, one-way ANOVA.

**Supplementary Fig. 4:**
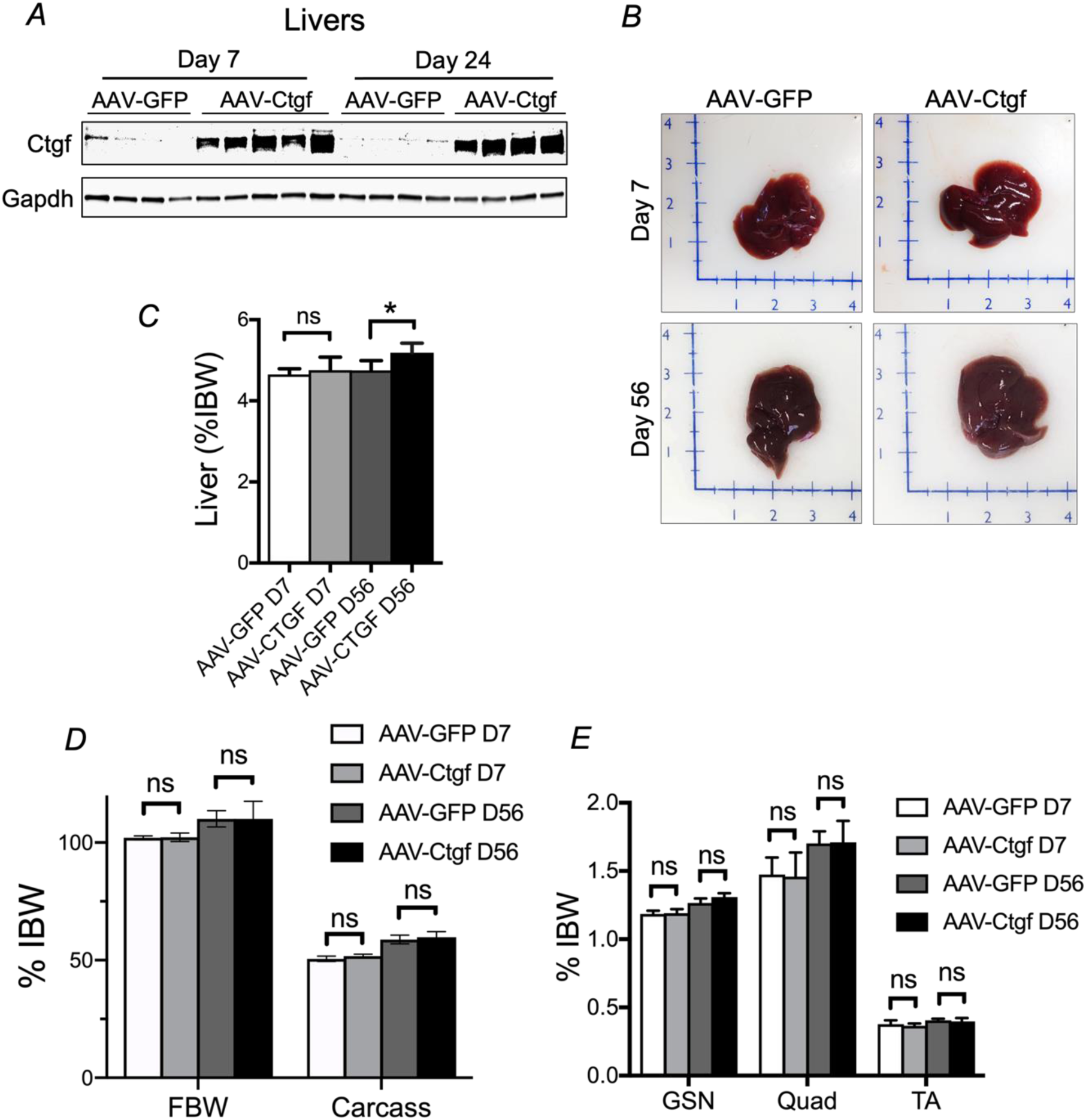
The Yap1 target gene *Gtgf* does not induce hepatomegaly and muscle loss in mice. A) Ctgf western blot in AAV-GFP and AAV-Ctgf livers, 7 and 24 days post-infection. B) Representative liver images in each AAV condition at day7 and day56 post-infection. C) Liver mass in each condition. AAV-GFP: n=4, AAV-Ctgf: n=5, \**P*<0.05, one-way ANOVA. D) Final body weight and carcass weight in each AAV condition. ns= not significant, one-way ANOVA. E) Skeletal muscle mass in each AAV condition. ns= not significant, one-way ANOVA.

**Supplementary Fig. 5:**
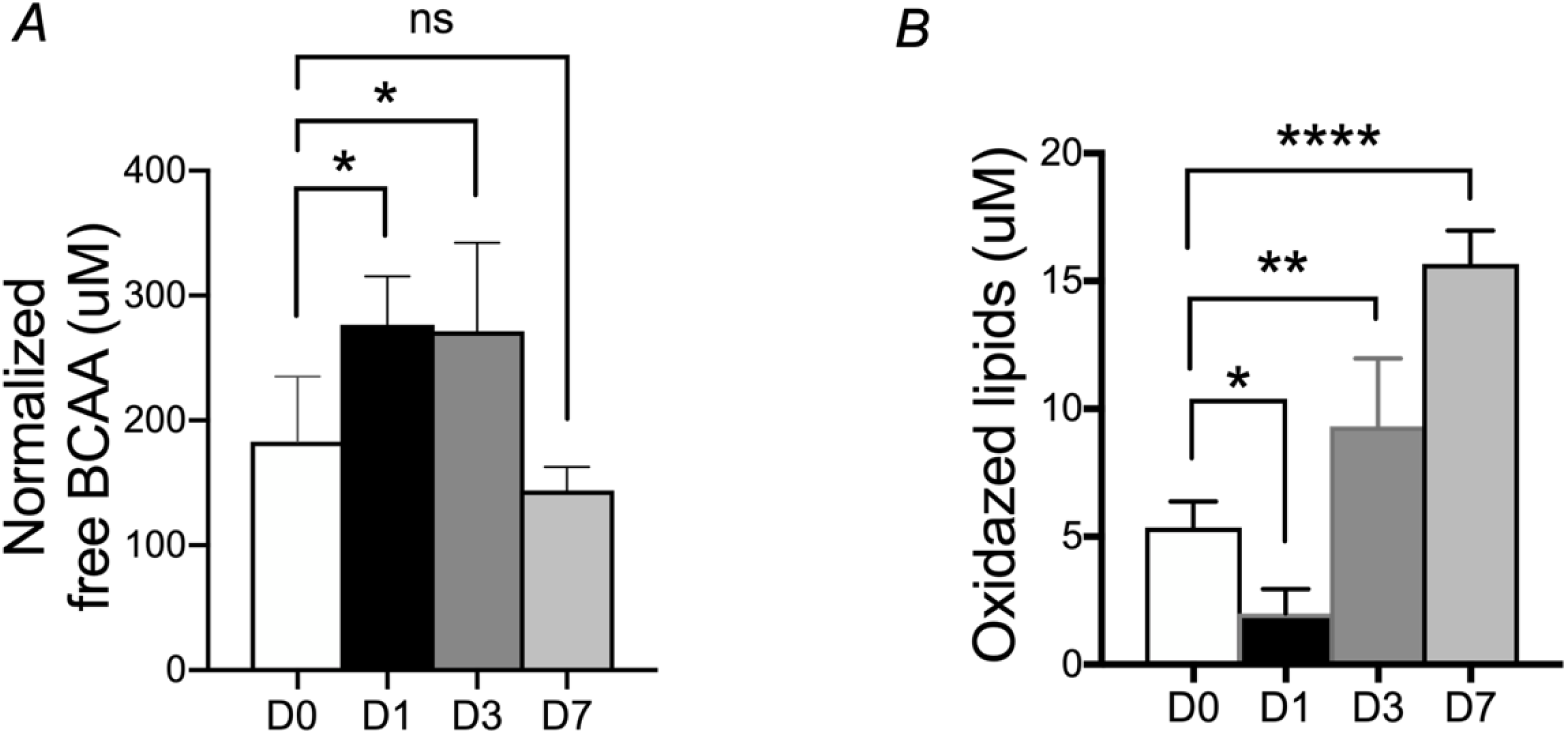
Elevated free BCAA and oxidized lipids in post-PHx blood plasma. A) Normalized free BCAA (uM) in blood plasma in d0, d1, d3 and d7 post-PHx. n=5 mice per time point. \**P*<0.05, one-way ANOVA. B) Oxidized lipids (uM) in blood plasma of d0, d1, d3, d7 post-PHx. n=5 mice per time point. \**P*<0.05, \*\**P*<0.01, \*\*\**P*<0.001, one-way ANOVA.

